# Phages weaponize their bacteria with biosynthetic gene clusters

**DOI:** 10.1101/2020.10.01.322628

**Authors:** Anna Dragoš, Aaron J.C. Andersen, Carlos N. Lozano-Andrade, Paul J. Kempen, Ákos T. Kovács, Mikael Lenz Strube

**Affiliations:** Department of Biotechnology and Biomedicine, Technical University of Denmark, Søltofts Plads bldg. 221, DK-2800 Kgs Lyngby, Denmark; Department of Health Technology, Technical University of Denmark, Produktionstorvet bldg. 423, DK-2800 Kgs Lyngby, Denmark

**Keywords:** Biosynthetic gene cluster, genome mining, phages, bacteriocins, interbacterial competition

## Abstract

Bacteria produce many different specialized metabolites, which are encoded by biosynthetic gene clusters (BGCs). Despite high industrial relevance owing to broad bioactive potential of these metabolites, their ecological roles remain largely unexplored. We analyze all available genomes for BGCs of phage origin. The BGCs predominantly reside within temperate phages infecting certain commensal and pathogenic bacteria. Nearly all phage BGCs encode bacteriocins, which appear to serve as a strong proxy for phage specificity. Using the gut-associated bacterium *Bacillus subtilis*, we demonstrate how a temperate phage equips its host with a functional BGC, providing it with a competitive fitness advantage over close relatives. Therefore, certain temperate phages use BGCs to weaponize their bacteria against close relatives, leading to evolutionary benefits from lysogeny to the infected host, and hence, to the phage itself. Our study is a large step towards understanding the natural role of specialized metabolites, as well as mutualistic phage-host relationships.

## Introduction

Apart from molecules directly involved in basic metabolism, microbes produce a plethora of specialized compounds, the so-called specialized metabolites. These metabolites mediate microbial ecological interactions, but also supply medical^1^ and biotech industries^2^ with novel biological activities. Despite a long tradition of industrial exploitation, the natural role of most specialized metabolites remains largely unknown^3^, although there is evidence that some of them act as chemical weapons in competitive interaction^4,5^.

Enzymatic pathways required to produce specialized metabolites are encoded by biosynthetic gene clusters (BGCs). Although BGCs differ both structurally and functionally, major groups with similar biosynthetic function are found, e.g. polyketide synthases (PKSs) and non-ribosomal peptide synthetases (NRPSs), which are extremely large (10-100 kb), multi-enzyme complexes synthesizing products with such properties as: antibiotic, signaling, immunosuppressive and biosurfactant activities^6,7^. Ribosomal peptide natural products (RPNPs or RiPPs) are mostly represented by bacteriocins, recognized for their role in interbacterial competition^8^. and their biosynthetic gene clusters are smaller compared to PKSs and NRPSs. The presence (and expression) of a BGC on a bacterial genome can determine its pathogenicity^9,10^, the potential for its success in an ecological niche^11^, and recently, we are becoming aware of the importance of specialized metabolites in human microbiomes in the balance between health and disease^12^. Regardless of the particular niche of a microorganism, it is evident that it will be locked in a perpetual state of chemical warfare with others^4^ and given the antibiotic activity of many specialized metabolites, we assume that specialized metabolites are crucial for the assembly of microbiomes. It has been proposed that stability in microbiomes is promoted when interactions are few and predominantly competitive^5^, suggesting that not only the individual bacteria, but the community as a whole benefits from hosting a BGC that encode an antagonistic compound(s). Bacteriocins are particularly interesting in this context as they are uniformly bactericidal in contrast to more complicated specialized metabolites which may have entirely divergent effects depending on the concentration^13^.

Progress in metagenomics along with excellent computational tools to predict BGCs^14^ have allowed us to characterize the genetic potential for production of specialized metabolites in microbiomes, as well as identify bacteria which are the main hosts of BGCs ^15–17^. For instance, in soil microbiomes, *Acidobacteria* tend to be the richest in BGCs, mainly synthesizing polyketides, non-ribosomal peptides or terpenes, however the *Actinobacteria* phylum contains multiple ‘outliers’ with extremely high (>20) BGC cargo^16^. Moreover, the abundance of BGCs of soil microbiomes also varies depending on vegetation and soil depth^16^. In the healthy human gut microbiome, *Bacteroides* genus serves as the major BGC-carrier, with saccharides being the most abundant specialized metabolites, while common gut-associated genera like *Escherichia, Lactobacillus, Haemophilus* or *Enterococcus*, appear limited in BGC cargo^15^. In the human oral microbiomes, the *Firmicutes* are the main carriers of BGCs, with oligosaccharide-encoding clusters and RiPPs as the major BGC types^12^. Although current interdisciplinary approaches allow us to identify the key BGC types and their producers, understanding the natural function of specialized metabolites and the eco-evolutionary factors behind their distribution in microbial communities remains challenging.

In certain bacterial genera, BGCs are commonly associated with mobile genetic elements (MGEs)^18^, the so-called flexible genome. For instance large BGCs are commonly found on plasmids of *Pseudonocardia*^18^, but whether, or how, being part of MGEs translate into structure and ecological function of the specialized metabolite remains unknown. This knowledge can be one of the large steps towards understanding the role of specialized metabolites in bacterial communities and possibly also their role in balance between health and dysbiosis^19^.

Amongst MGEs, phages are considered the most abundant biological agents on Earth. Phages have a major impact on bacterial biomass^20^ and community composition, such as serving as important stimulus of healthy gut microbiota^21^. Lytic phages hijack the DNA and protein synthesis machinery of the host, rapidly multiply and eventually lyse the host cell to release progeny virions. Temperate phages can integrate their DNA into the host chromosome using specific or unspecific attachment sites (*att*) and replicate with the host bacterium as a prophage. Most bacterial genomes carry at least one prophage^22^, and prophage elements can occupy up to 20% of the entire bacterial genome^23^.

Phages can encode virulence factors^24,25^, toxins^26^ and other compounds potentially valuable for the host. For instance, the SPβ prophage present in some strains of a soil, plant and gut bacterium *Bacillus subtilis*^28^, encodes for an S-linked glycocin called sublancin^29,30^. The sublancin cluster (5 kb) contains a precursor peptide (*sunA*), posttranslational modification enzymes (*sunS, bdbA, bdbB*), a self-immunity protein (*sunI*), and a transporter (*sunT*) that ensures the export of the bacteriocin into extracellular space^31^. Sublancin presumably enters the cells using the PTS-glucose specific transport system^32^, and upon entry, negatively affects DNA replication, transcription and RNA translation^33^. Mutants deprived of SPβ, the entire sublancin cluster or just the immunity gene *sunI*, are inhibited by isogenic strains that carry intact SPβ^34^. This prophage can be found in closely related but geographically dispersed *B. subtilis* isolates, always occupying the same position in the chromosome^28^. The contribution of sublancin to ecological success of SPβ remains an open question. In the same line of thought, we do not know if other phages/prophages carry BGCs and whether the products of these BGCs may share structural or functional features. Such knowledge would contribute to our understanding of the natural role of bacterial BGCs, and would also facilitate mining and utilization of new specialized metabolites.

The aim this study was to explore all high-quality phage and prophage genomes for the presence of BGCs using a large-scale bioinformatics approach. We demonstrate that BGCs are more likely to be found within genomes of temperate phages which infect commensal/pathogenic bacterial hosts. The study also shows that nearly all phage-BGCs encode bacteriocins, suggesting they may serve as chemical weapons provided by phages to their bacterial hosts, hence increasing benefits from lysogeny. This hypothesis is experimentally confirmed, revealing the competitive benefits from bacteriocin-carrying phage acquisition by a natural *B. subtilis* isolate. Comparison of host phylogeny, with phage and BGC relatedness shows that bacteriocins serve as a good indicator of phage host range. Our work contributes significantly to understanding of ecological role of specialized metabolites, as well as cooperative relationship between phages and their bacteria.

## Results

### Biosynthetic gene clusters (BGCs) are extremely rare within virion-derived genomes

To assess the contribution of phages to the genetic pool of microbial specialized metabolites, we began from a basic question: can biosynthetic gene clusters (BGCs) that encode for specialized metabolites be found on phage genomes? To answer this question, all complete phage genomes available in the PATRIC 3.6.2 database (10063 in total) were subjected to BGC detection by antiSMASH 4 (see methods). Interestingly, only approximately 0.07 % of all phages (69 in total) carried BGCs (Fig. 1A). These phage-encoded BGCs will be referred to as pBGCs (phage-encoded biosynthetic gene clusters).

**Fig. 1.**
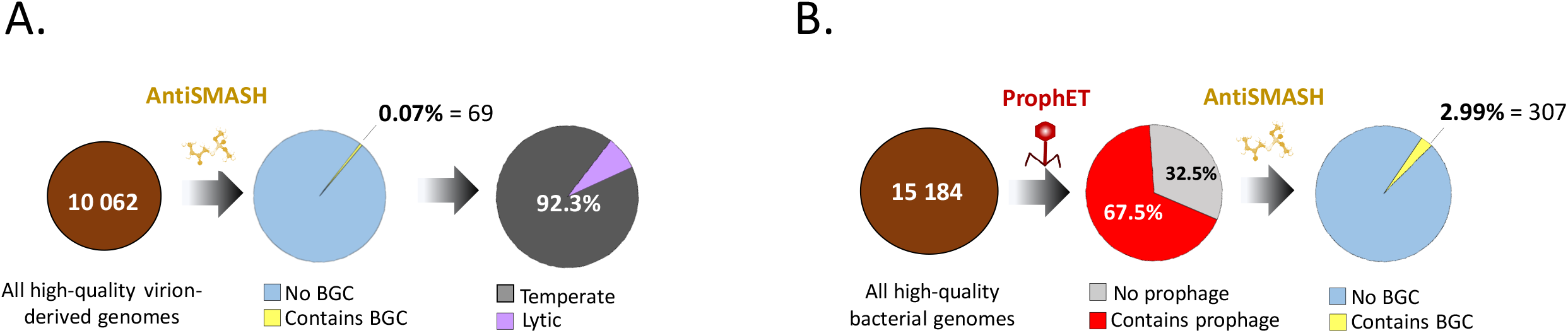
Bioinformatic worflow used to extract phage- or prophage-encoded biosythetic gene clusters (pBGCs). A. All high-quality phage genomes available in PATRIC database (10062) were screened by antiSMASH, and only 0.07 % appeared to contain pBGCs. The vast majority of these pBGCs-containing phages (92.3 %) were capable of lysogenic lifestyle, as confirmed by literature mining. B. All high quality bacterial genomes available from NCBI (15184) were screened by ProphET to extract prophage rergions, which were next screened by antiSMASH to detect pBGCs. Relative abundance of pBGCs within prophages was nearly 3 %.

The vast majority (64) of pBGCs were found in temperate phages (for which lifestyle was confirmed experimentally in the literature) and only five pBGCs were carried by a lytic phage (life style confirmed experimentally in the literature) (Suppl. dataset 1). We also noticed that phages infecting certain bacterial genera were overrepresented among the pBGC hits, compared to relative distribution of phage hosts in PATRIC database (Suppl. dataset 1, Suppl. Fig. 1A). These genera where *Escherichia* (represented solely by *Escherichia coli* species), *Mannheimia, Enterobacter* and *Shigella* (Fig. 2A, Suppl. Fig. 1A).

**Fig. 2:**
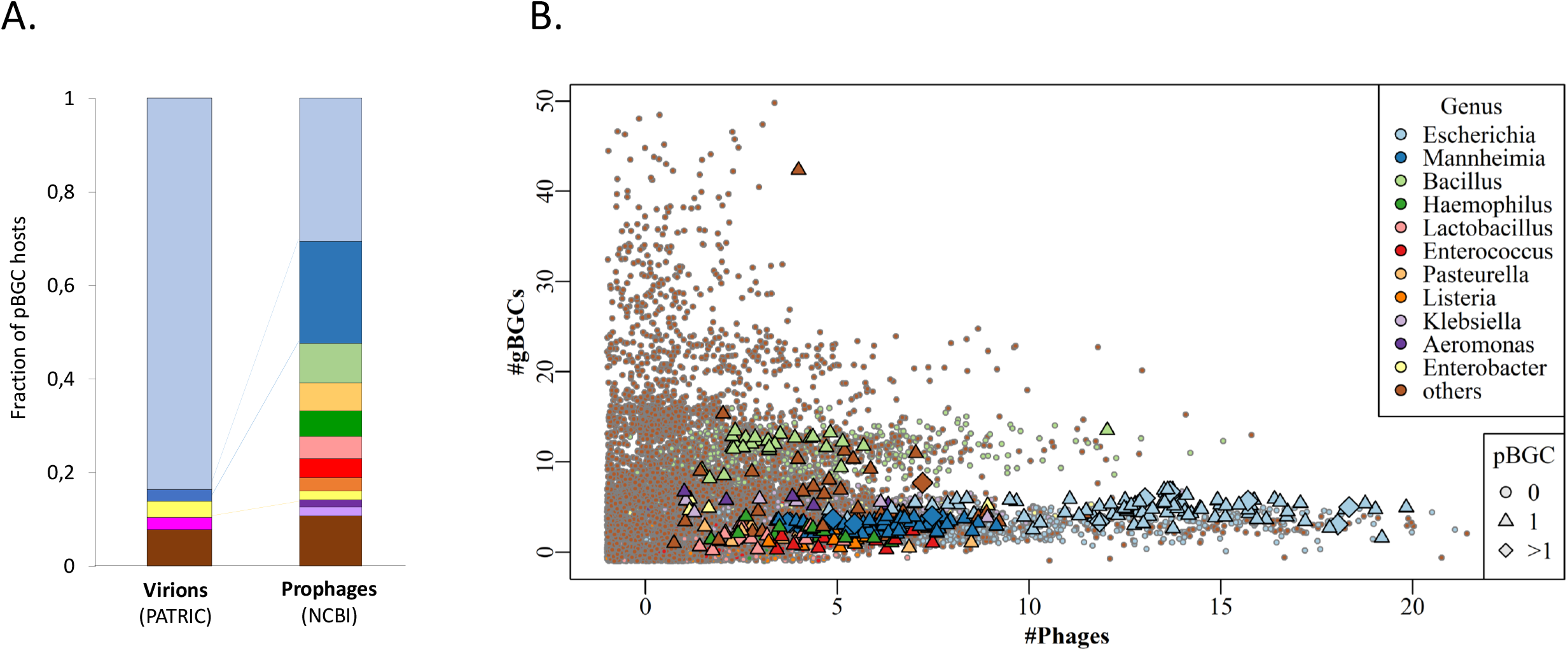
Distribution of pBGC across genera. A) Relative distribution of bacteria which serve as hosts to pBGC-carrying phages colored by genus. For colors interpretation, see figure legend in 2B. Pink indicates Shigella genus. B) Number of prophages in each genome versus number of genomic BGCs colored by genus. Genomes having only one pBGC are marked by triangles and genomes containing multiple pBGCs are marked by diamonds. Points have been jittered to avoid overlap.

It is important to note that number of BGC hits per phage genome was restricted to 1 or none in all cases. This initial analysis suggested that pBGCs are extremely rare, and that they are more likely to reside within host-associated (temperate) phages infecting particular bacterial species.

### BGCs are more abundant in prophage elements compared to virions

Since the previous analysis suggested that pBGCs may be more common within temperate phages, the pBGCs encoded by prophage elements were investigated. As the pBGC-positive phage pool is largely overrepresented by *Escherichia coli* phages (Suppl. dataset 1), the aim was to analyze a more diverse set of (pro)phage genomes. Prophage regions were predicted and extracted from all high-quality bacterial genomes available in the NCBI database, and subjected these to BGC detection by antiSMASH 4 (see methods) (Fig. 1B). We found that the majority of analyzed bacterial genomes (67.5 % out of 15184 total genomes analyzed) carried at least one prophage element, and only approximately 3 % of these prophage regions (307 in total) contained a BGC (Fig. 1B, Suppl. dataset 2) corresponding to approximately 2 % of all bacterial genomes.

It was again observed that certain host species (including three out of four that were also ‘popular’ host genera of pBGC-carrying virions) were overrepresented among pBGC-carrying prophage elements compared to relative abundance of these genera in the entire genome database (Suppl. dataset 2, Suppl. Fig 1B). Specifically, nearly 90 % of all pBGC-carrying prophages were associated with 11 genera of human/animal commensals and pathogens: *Escherichia* (in which all but 1 isolate belonged to *E. coli*), *Mannheimia* (where all isolates belonged to *Mannhemia heamolytica*), *Bacillus, Pasteurella, Haemophilus, Lactobacillus, Enterococcus, Enterobacter, Listeria, Aeromonas* and *Klebsiella*. All, apart from the latter two, seemed to be several-fold overrepresented within our pBGC dataset (Fig. 2A, Suppl. dataset 2, Suppl. Fig 1B). However, not all representatives of these genera had pBGCs. In addition, pBGC were clearly over-represented in prophages of certain species, e.g. *B. subtilis* which is in the minority (20 %) of all *Bacillus* genomes in NCBI, but it makes up for the majority (74 %) of pBGC hits within *Bacillus* genus; *Lactobacillus brevis* (4 % of all lactobacilli but majority (67 %) of pBGC hits within *Lactobacillus* genus), or *Enterococcus faecalis* (22 % of all enterococci and over 90 % of pBGC hits within corresponding genus) (Suppl. dataset 2, Suppl. Fig 1B).

In the vast majority of pBGC-carrying bacteria, the prophage carried only one BGCs, with the notable exceptions being some isolates of *M. haemolytica* (8 isolates), *E. coli* (7 isolates) and *L. brevis* (1 isolate) carrying 2 pBGCs. A single genome belonging to *Alkaliphilus metalliredigens* carried three pBGCs, all of which were identical (Supp. dataset 2).

Since total number of phages were substantially higher in pBGC hosts (Fig. 2B, Mann-Whitney U-test, P=2.2*10^−16^). We then asked whether prophages with BGC cargo are simply more abundant within species that are generally rich in BGCs. The number of BGCs encoded outside the prophage regions (which will be referred to as gBGC for core-genome-encoded biosynthetic gene cluster) were assessed (Suppl. dataset 2) revealing the opposite scenario: total gBGC count was slightly, but significantly, lower in pBGC hosts (Fig 2B, Mann-Whitney U-test, P=0.0011).

Consequently, in certain species like *Listeria monocytogenes, E. faecalis*, or *L. brevis* which have low numbers of gBGCs, phages appeared as a major or even a sole source of biosynthetic gene clusters (Fig. 2). These results confirm that, although relatively rare, pBGCs are more likely to be found within temperate phages, infecting certain species of human/animal commensals or pathogens, with limited gBGC cargo.

### The vast majority of prophage-encoded BGCs are bacteriocins

We then investigated if prophage-encoded BGCs share structural/functional features. Indeed, according to antiSMASH prediction, nearly all pBGC were assigned to bacteriocins (99.3 %), with some stratification onto glycocins (n=22, 6.8 %) and lantipeptides (n=2, 0.6 %). To investigate how many of the antiSMASH-predicted pBGC belonged to experimentally confirmed/studied bacteriocins, the pBGCs were compared with the BACTIBASE database (see methods). The vast majority of pBGCs had poor matches in BACTIBASE, most having protein similarities and alignment lengths well below 60 % or lower (Fig. 3). Examples of pBGCs of high similarity to experimentally confirmed bacteriocins were: sublancin (22 hits in *B. subtilis*), enterocin (5 hits in *L. monocytogenes*) and carocin (*Pectobactrerium carotovorum* (GCF_000294535.1)). Linocin was found in a close to full length, but highly dissimilar, version in *Nocardia terpenica* (GCF_002568625.1) and *E. coli* (GCF_001901005.1 and GCF_900636075.1).

**Fig. 3:**
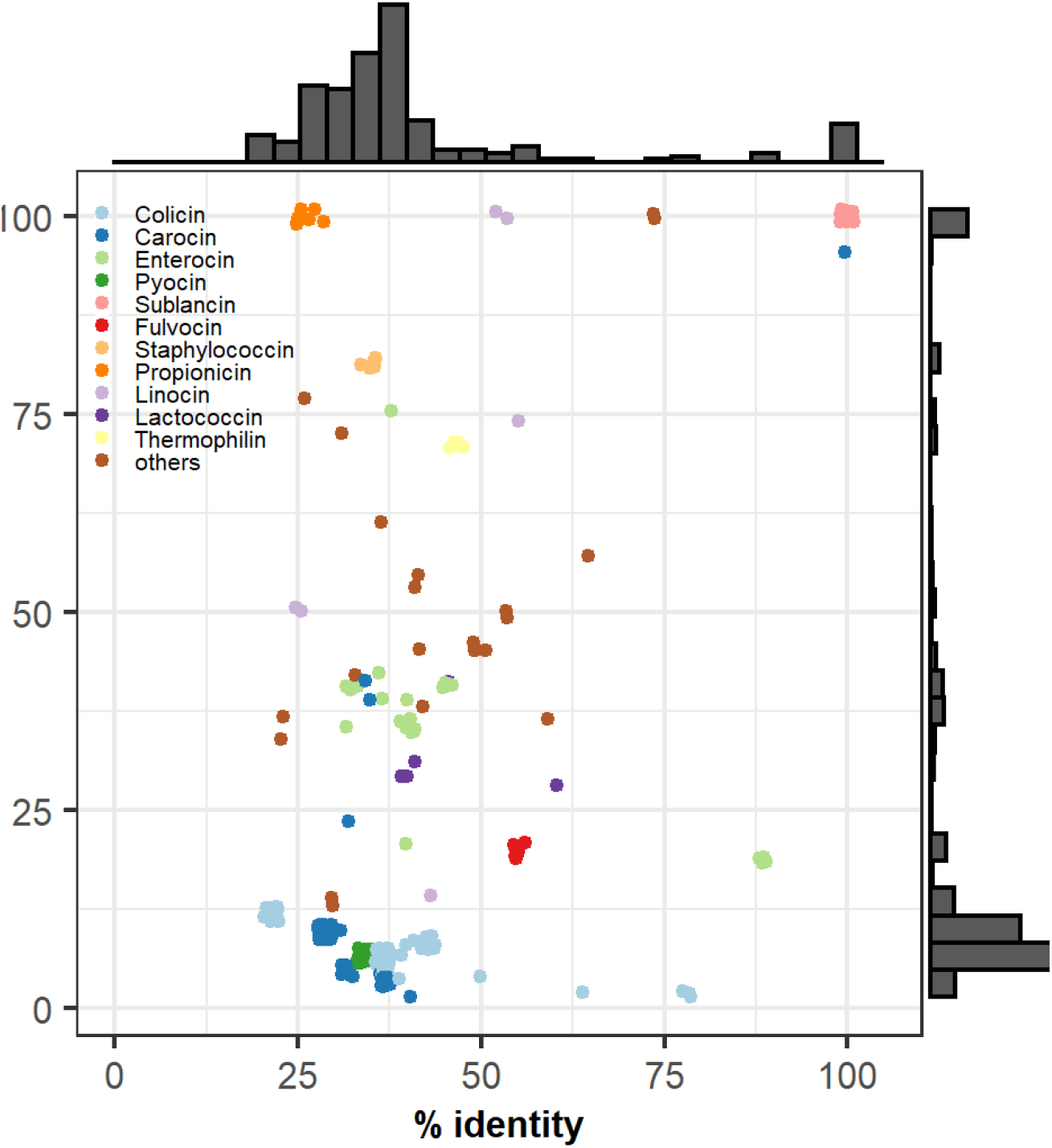
BLASTP identities and alignment lengths of pBGC core proteins. Points are colored by their closest match and have been jittered to avoid overlap. Histograms show densities for the corresponding blastp metric.

Although the vast majority of pBGCs were bacteriocins, we have also found several exceptions. For instance, 2 pBGCs found in two separate *Pseudomonas aeruginosa* genomes contained a partial NRPS-like cluster with a single AMP-binding domain as well as a siderophore receptor and a sigma-70 polymerase sigma factor (Suppl. dataset 2). Two pBGCs found within 2 *Mycobacterium* temperate phages encoded for ectoine which may play a role in osmoadaptation, a pBGC found on the pseudo-lysogenic phage *Streptomyces* phage ZL12 was annotated as a Type 3 PKS, while a pBGC found in a lytic *Roseobacter* phage encoded for homo-serine lactone, which is a molecule involved in cell-to-cell communication (Suppl. dataset 1).

These results suggest that phages can be the source of different BGC types, however the vast majority of pBGCs are bacteriocins – putative weapons of competition against closely related bacteria.

### Phages weaponize their hosts with BGCs through lysogeny

Since pBGCs are more commonly found in temperate phages, it was hypothesized they may play an important role during lysogeny. As the vast majority of pBGCs are bacteriocins, they could serve as biological weapons during competition of the host bacterium with its closely-related neighbors, thereby increasing fitness benefits from lysogeny for both host and phage. This hypothesis was tested using a *B. subtilis* model system, by transferring a pBGC carrying phage from one natural isolate to another and monitoring fitness benefits from such pBGC-phage infection (Fig. 4A). The isolate MB8_B7 (CP045821)^35^ served as a phage donor, as it is a natural host of an SPβ phage, which carries the biosynthetic gene cluster of the antimicrobial glycocin, sublancin (Fig. 4A, Suppl. Fig. 2A). The pBGC-negative isolate P9_B1 was used as a receiver strain lacking SPβ phage in the genome (CP045811)^35^.

**Fig. 4.**
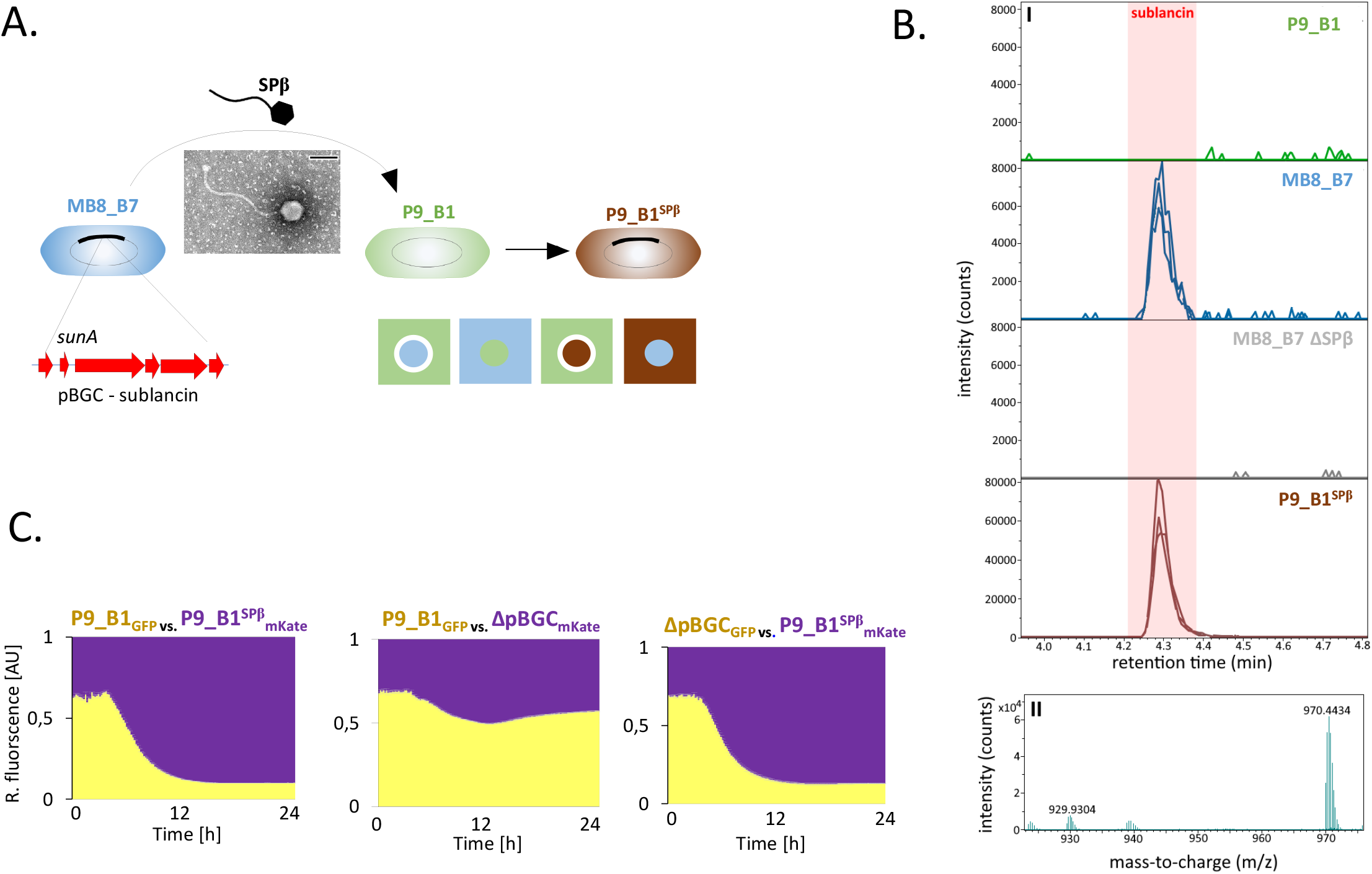
Phages use pBGC to weaponize their bacterial host. A. *B. subtilis* isolate MB8_B7 (pictured in blue) is lysogenic for the SPβ prophage which carries a pBGC responsible for sublancin biosynthesis, *sunA* being the core gene. The SPβ prophage was induced and purified, as confirmed by transmission electron microscopy (scale bar = 100 nm) and used to infect the soil isolate P9_B1 (pictured in green) resulting in P9_B1^SPβ^ (pictured in brown). Antagonistic activity/immunity of MB8_B7, P9_B1 and P9_B1^SPβ^ were indicated as square diagram, where square represents a lawn and dot represents a focal strain. A white ring around the focal strain indicates its antagonistic activity against the lawn strain. B. Detection of sublancin in supernatants of P9_B1, MB8_B7, MB8_B7 ΔSPβ and P9_B1^SPβ^ using LC-MS. C. Competition between P9_B1_GFP_ vs. P9_B1^SPβ^_mKate_; P9_B1_GFP_ vs. ΔpBGC_mKate_ and ΔpBGC_GFP_ vs. P9_B1^SPβ^_mKate_ starting from 1:1 inoculation ratio. Relative abundance of two competing strains was monitored by changes in relative GFP and mKate fluorescence, shown in yellow and purple, respectively. Error bars (light grey) indicates standard error (n=6).

The genome sequence of the MB8_B7 prophage was identical to the SPβ-phage found in the lab strain *B. subtilis* 168, except that the MB8_B7 does not contain ICEBs1, shown to interfere with the lytic cycle of SPβ^36^ (see methods). It was confirmed that deletion of SPβ from MB8_B7 provoked sensitivity towards the wild-type (WT) MB8_B7 (Fig. 4A, Suppl. Fig. 2B). Importantly, sublancin could be detected in the spent medium of MB8_B7 but not in its ΔSPβ derivative (Fig. 4B).

Next, SPβ virions were isolated from MB8_B7 culture (see methods), confirmed its identity by Nanopore sequencing (see methods) (Suppl. Fig. 6A) and used it to infect the naturally pBGC-free isolate P9_B1 (Fig. 4A, Suppl. Fig. 2CD). As anticipated, the P9_B1^SPβ^ became antagonistic towards its P9_B1 ancestor (Fig. Suppl. Fig. 2C). The P9_B1^SPβ^ was also able to produce sublancin (Fig. 4AB). These results clearly demonstrated that the phage-encoded BGC and its biological activity can be easily transferred between two strains via phage infection (Fig. 4AB).

**Fig. 6.**
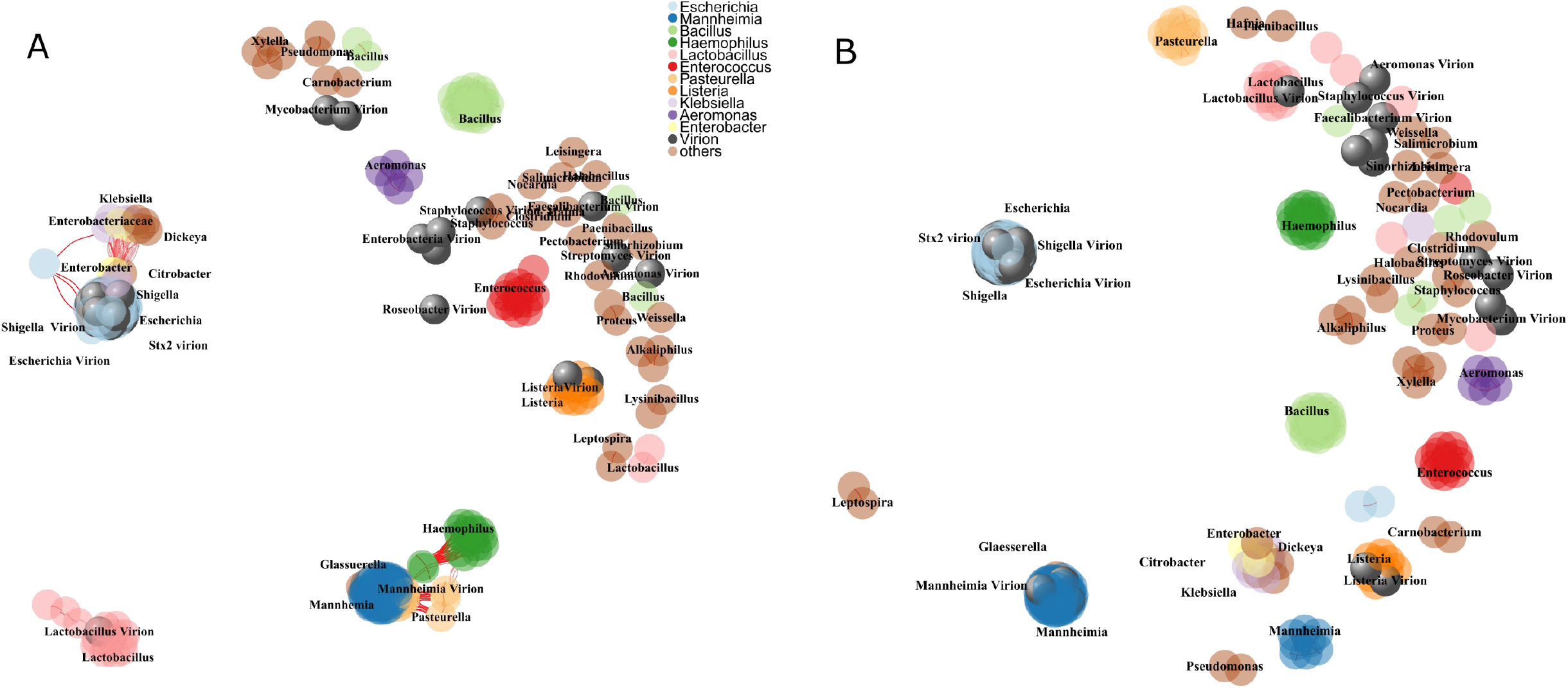
Diversity of phages and pBGCs. A) Network of pBGC similarity of full length prophages and virions carrying pBGCs, labelled by major genera. Each edge is weighted by the ANI-value of each pair. B) Network of pBGC core gene similarity of prophage- and virion-derived pBGCs, labelled by major genera. Each edge is weighted by the ANI-value of each pair.

To further examine benefits from pBGC acquisition, the P9_B1 and P9_B1^SPβ^ were labelled with fluorescent reporters (see methods; Suppl. Fig. 3A), and followed their growth in co-cultures, starting from 1:1 ratio.

Clearly, P9_B1^SPβ^ was winning over P9_B1, regardless of the fluorescent marker used (Student’s *t*-test, P= 0.694 ×10^−13^, P= 0.160 ×10^−7^, for P9_B1_GFP_ vs. P9_B1^SPβ^_mKate_ and P9_B1_mKate_ vs. P9_B1^SPβ^_mKate_ respectively, see methods) (Fig. 4C, Suppl. Fig. 4A). To confirm that competitive advantage of P9_B1^SPβ^ was due to pBGC (sublancin), we deleted the entire pBGC (sublancin production as well as immunity gene) or only the core gene *sunA* in P9_B1^SPβ^ and challenged the mutants against the P9_B1 or P9_B1^SPβ^. Both ΔpBGC and Δ*sunA* strains lost their competitive advantage over the P9_B1 (Fig. 4C, Suppl. Fig. 4AB). In addition, ΔpBGC, but not Δ*sunA* was now loosing against P9_B1^SPβ^ most likely due to lack of immunity to sublancin (Student’s *t*-test, P= 0.664 ×10^−10^, P= 0.301 ×10^−17^, for P9_B1^SPβ^_GFP_ vs. ΔpBGC_mKate_ and P9_B1^SPβ^_mKate_ vs. ΔpBGC_GFP_ respectively, see methods) (Fig. 4C, Suppl. Fig. 4B). As a control for potential fitness effects of the fluorescent markers, the co-cultures of isogenic strains were also examined, but the ratios remained as inoculated (Suppl. Fig. 4C).

We took an alternative approach to examine competitive advantage from carrying an attachment site (*att*) for pBGC phage on the chromosome. As not all *B. subtilis* strains carry *spsM*^28^, the lack of SPβ attachment gene would greatly reduce the frequency of SPβ integration into the host chromosome (Fig. 5A), and in effect, render the *spsM*-mutant less competitively fit compared to the wild type. We designed a subsequent inoculation-based competition assay with P9_B1 vs. Δ*spsM* in absence of SPβ (control) and with a single episode of exposure to SPβ (see methods) (Fig. 5B). Both strains showed similar susceptibility to lysis by SPβ, based on growth curves and plaque assay (Suppl. Fig. 5A).

**Fig. 5.**
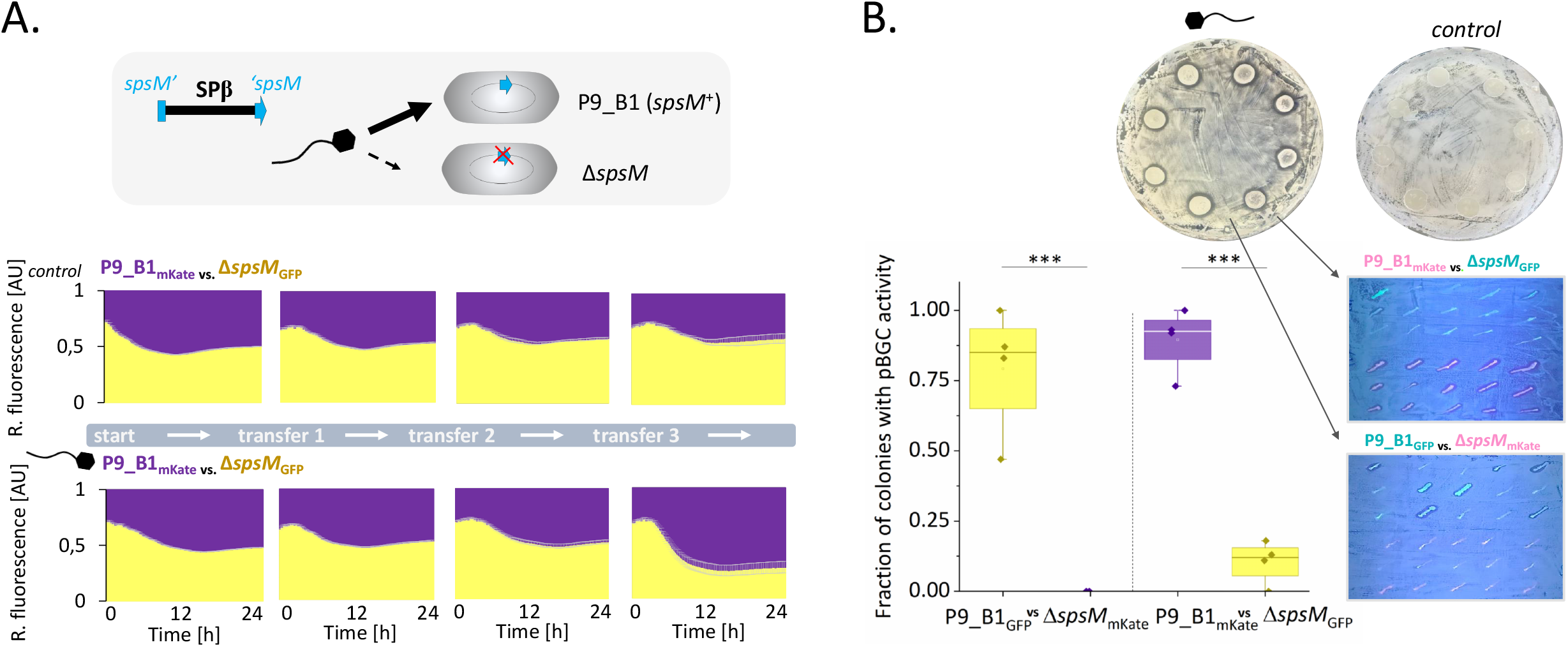
Bacteria benefit from attachment gene for pBGC-carrying phages. A. Graphics above – SPβ typically integrates into *spsM* gene splitting this gene. It was hypothesized that frequency of lysogenic conversion will be greatly reduced if *spsM* gene is removed. Below – long term competition assay between initially SPβ-negative P9_B1_mKate_ vs Δs*psM*_GFP_ without and with single exposure to SPβ at the exponential growth phase during first co-cultivation round. All assays were initiated at 1:1 proportions. Relative abundance of two competing strains was monitored by changes in relative GFP and mKate fluorescence, shown in yellow and purple, respectively. Error bars (light grey) indicate standard error (n=8). B. Mixed populations of P9_B1 and Δ*spsM* after long-term cultivation experiment were screened for antagonistic activity against the P9_B1 ancestor. All co-cultures exposed to SPβ acquired antagonistic activity towards the P9_B1, which was not observed in any of the control co-cultures (no phage solution added). P9_B1 and Δ*spsM* were separated to single colonies and again screened for antagonistic activity against the P9_B1 ancestor by striking the colonies on top of P9_B1 lawn. Plates were illuminated by UV lamp and imaged by iPhone7 and the fraction of P9_B1 and Δ*spsM* colonies inhibitory towards P9_B1 was calculated. Green strikes indicate GFP-labelled strains and pink strikes indicate mKate-labelled strains. *** indicates P<0.001 (n=4).

In the absence of the phage, both strains could co-exist at comparable relative proportions for at least 4 rounds of cultivation (see methods) (Fig. 5A, Suppl. Fig. 5B). However, a single exposure to SPβ at the beginning of the experiment dramatically changed the competition outcome, resulting in higher frequency of the P9_B1, regardless of fluorescent reporter used (p< 0.125×10^−3^, p< 0.423 ×10^−3^ for P9_B1_GFP_ vs. Δ*spsM*_mKate_ and P9_B1_mKate_ vs. Δ*spsM*_GFP_ respectively, see methods) (Fig. 5A, Suppl. Fig. 5B). To test whether the P9_B1 was winning due to higher rate of lysogenic conversion and pBGC acquisition, the strains were separated to single colonies and tested their antagonistic activity against the P9_B1 ancestor (Fig. 5B). Indeed, the vast majority of P9_B1 colonies were antagonistic against the ancestor, which was not the case for Δ*spsM*. Finally, PCR screening of 4 randomly selected colonies from the Δ*spsM* samples confirmed that the presence of antagonistic activity correlated with integration of SPβ-specific regions (including sublancin) into the chromosome (Suppl. Fig. 5C). Using Nanopore sequencing, SPβ integration was confirmed in an alternative locus (*yxzL*) in absence of main *att* site in the bacterial chromosome (Suppl. Fig. 6B).

These results demonstrated that a phages-borne pBGC confers a fitness benefit to the host bacterium, in turn provokes a fitness benefit for the phage spreading as well. In addition, bacteria benefit from carrying attachment sites in their genomes where pBGC-carrying phages can integrate.

### pBGCs-carrying phages are more promiscuous than the pBGCs they carry

We then assessed the role of phages in horizontal gene transfer (HGT) of BGCs between closely and more distantly related bacteria. First, all pBGCs prophages and virions along with their corresponding pBGCs core genes were compared by average nucleotide identities. This allowed assessment of pBGC conservation level within host genera, as well as the potential of phages to transfer pBGCs across genera and family. When comparing core pBGC genes by average nucleotide identity, clustering according to bacterial host genera was evident (Fig. 6, Suppl. Fig. 7). Most pBGC carrying virions retrieved from the PATRIC database clustered according to their PATRIC-annotated hosts and with ProphET-identified prophages found in the corresponding host genus (Fig. 6, Suppl. Fig. 7). This confirms presence of a functional lytic cycle in some pBGC-carrying prophages targeting *Escherichia, Mainhemia, Listeria* and *Lactobacillus* opening up the possibility of pBGCs transfer via phage infection in nature.

Comparing the sequence similarity of pBGC carrying phages with the similarity of core pBGC genes, we could identify 4 distinct patterns: 1) phage relatedness matches pBGCs relatedness; 2) similar pBGCs can be found on distinct phages; 3) distinct pBGCs can be found on similar phages; 4) pBGC can be shared across genera. First group is represented by *Bacillus, Lactobacillus*, or *Listeria*, where pBGCs carrying phages as well as the pBGC core genes clearly clustered together (Fig. 6, Suppl. Fig. 7). For *Listeria*, the pBGCs carrying phages, and pBGC core genes are nearly identical within the genus. Although the remaining genera (*Bacillus* or *Lactobacillus*) contain some outlier pBGCs, which do not belong to the main pBGC clusters, these were always associated with outlier prophages found within the corresponding genera. This suggests that in *Bacillus, Lactobacillus* and *Listeria*, potential horizontal gene transfer of pBGC may be restricted to the genus level.

The second group is represented by *Mainhemia*, where all pBGCs are identical within genus, but they can be found on diverse prophage elements (Fig. 6, Suppl. Fig. 7). This either indicates that pBGCs of *Mainhemia* undergo between-phage horizontal gene transfer, or that they remain under purifying selection– for instance due to their beneficial function for the bacterial host (Fig. 6, Suppl. Fig. 7).

The third group is represented by *Enterococcus*, where prophage diversity is rather limited (Fig. 6, Suppl. Fig. 7) but the pBGC core genes are clearly divided into two different clusters (Fig. 6, Suppl. Fig. 7). Similar phenomena can be observed even across different genera, like *Mainhemia, Pasteurella* and *Heamophilus*, where the latter two carry *Mainhemia*-like prophages, but the pBGCs core genes remain unique for each genus (Fig. 6). The same is the case for *Aeromonas* and *Klebsiella*, which carry *Escherichia*-like prophages, but not *Escherichia*-like pBGCs core genes (Fig. 6), Suppl. Fig. 7. This may indicate that pBGCs play a role in early diversification of phages towards different host species/genera. BLAST analysis confirmed that most pBGCs (290 out of 307) had a match (>70 % id, >70 % length) within genome of another pBGC carrier and only 17 were unique pBGCs as also observed from ANI plots (Fig. 6AB, Suppl. Fig. 7).

BLAST comparison also revealed that 19 pBGC carriers hosted an additional copy of a highly similar (80-90 %) bacteriocin outside the original prophage region (Suppl. dataset 3). We also found that 5.1 % of genomes which were originally identified as non-pBGC carriers (14877 in total Fig. 1B) had at least one analog to the pBGC core genes: in 56.4 % of these, pBGC analogues were found in the same species as the original pBGC host, 38.0 % homologues were found in different species within the same genus as the pBGC host, while 7.3 % were found in different genera, but the same family (e.g. *Escherichia* > *Shigella, Manheimia* > *Glaesserella*).

These results indicate that despite promiscuity of certain pBGCs carrying phages, and high potential for cross-genus transfer of pBGCs, pBGCs core genes are rarely shared between different host genera.

### Discussion

Specialized metabolites play an important role in microbial interactions and are exploited by the pharmaceutical industry, however, understanding the role and purpose of these compounds in nature has only recently become an area of research. Here, we explored a novel concept suggesting that phages may function as vectors for transferring biosynthetic potential for specialized metabolites therefore contributing to their natural distribution and ecological impact. Although BGCs are rarely found within phage genomes, when they are, they mostly reside within temperate phages or prophages.

Our bioinformatics approach has several drawbacks inherent in the scale of our analysis. ProphET algorithm^37^ was used for prophage recognition as it is currently the best-performing stand-alone prophage finder, although the reliance on a prophage database may underestimate novel prophages. AntiSMASH^38^ is currently the gold standard for profiling genomes for all types of BGCs, including genes encoding bacteriocins.

Considering the relatively high abundance of other phage-encoded secreted compounds (e.g. toxins and other virulence factors) which impact bacterial ecology^25,39,40^, one could expect phages to carry BGCs encoding for bacteriocins. On the other hand, hosting large multi-gene assemblies like BGCs is probably constrained by the efficiency of the phage DNA packing process, which translates to virion stability^41^. Such size constrains could also explain why phages accommodate the smallest BGCs – those encoding bacteriocins, which rarely exceed 10 kb^42^.

It was previously shown that host-associated lifestyle tends to correlate with higher prophage cargo^43^, which is in line with our findings on higher abundance of pBGCs in commensal/pathogenic species (Fig. 2A).

Host lifestyle cannot be a sole predictor for pBGC, since some of the recurrent facultative pathogens e.g. *Acinetobacter, Burkholderia, Pseudomonas* or *Staphylococcus*, contained no, or very few pBGCs, despite presence of prophages^44–46^ In addition, in the vast majority of polylysogens (strains infected with multiple phages) containing even up to 19 prophages (Suppl. dataset 2), there was only only pBGC-carrying prophage. This might be simply due to immunity of the pBGC-carrier to infection by another pBGC-carrying phage, since pBGC-carriers targeting the same species are likely phylogenetically related.

The fact that pBGCs were underrepresented in bacteria with high numbers of BGCs in the genome (gBGCs), could suggest that the presence of gBGC reduces the likelihood of phage infections, matching previous data on role of gBGC in anti-phage defense^47^. Alternatively, lysogeny could reduce the likelihood of gBGC acquisition, for instance by negatively influencing genetic competence for transformation^48,49^. Nonetheless, our results are in line with BGCs-survey within human microbiome project, where the leading pBGCs-carriers *Escherichia, Lactobacillus, Haemophilus* and *Enterococcus*,were identified to have lowest numbers of BGCs (less than 2)^15^.

Our work supports previous findings on temperate phages as carriers of putative weapons in inter-bacterial competition^26^. Previously, it was demonstrated that phages can repair BGCs by transducing missing genes between the strains^50^. Here, a complete and functional BGC is gained by prophage integration. Although the prophage itself (without pBGC) could serve as a biological weapon through spontaneous induction of lytic cycle in a subpopulation of cells^51^, this work demonstrates, in line with previous findings, that sublancin itself is sufficient to explain fitness benefits of lysogeny. The host also benefits from the intact phage attachment gene in presence of the pBGC phage. The *spsM* gene has previously been shown to be involved in biofilm production and sporulation, indicating that the absence of pBGC prophage can also serve competitive advantage under certain ecological conditions e.g. in the absence of close relatives, susceptible to sublancin attack ^52,53^. Here, the *spsM* locus provides function-independent benefits, uncovering fitness gain from certain chromosomal genes to capture pBGC phages more efficiently.

Recently, so called MuF polymorphic toxins have been predicted within multiple prophages of Firmicutes and proposed to serve as weapons in microbial warfare^26^. We see only minimal overlap between MuF toxin-carriers and pBGC-carrying phages from our dataset (9 bacterial genomes and only 5 prophages overlap) (compare Suppl. dataset 1 and Dataset S6^26^), confirming differences in genetic architecture, but not excluding similar ecological function, which in case of MuF toxins requires experimental validation. It is also important to distinguish between phage-encoded bacteriocins and well-described virulence factors residing in prophage regions e.g. the Shiga toxin encoded by *E. coli* lambdoid prophages, cholera toxin carried by *Vibrio cholerae* CTX phage or diphtheria toxin in *Corynebacterium diphtheriae* prophage, which target eukaryotic cells and may even hinder the inter-bacterial competition^54^. Discovering that nearly all pBGCs are bacteriocins also creates a fascinating connection to tailocins; bacterocidal phage tail-like particles, produced by certain genera, e.g. *Pseudomonas, Vibrio, Burkholderia* or *Clostridium*^56^. Although tailocin-carriers do not overlap with pBGC-carrying species, they probably share similar ecological role.

We observed that pBGC core genes match host phylogeny more than the entire phage genomes, which is in line with previous results comparing phage genome vs single protein networks with host phylogeny^57^. Previous work has revealed that the best phylogeny predictors are proteins that play a role in host recognition^57^. Could bacteriocins be a good predictor of host range and what is the potential mechanism? As phage specificity depends on recognition of specific receptors on the surface of host cells, by so called receptor binding proteins (RBPs), phage-encoded bacteriocins could recognize the same receptors. In fact, PTS-glucose specific transport system, a putative sublancin target^32^ is also known to play a role in phage entry in *E*.*coli* ^58^.

Finally, clustering of pBGCs according to genus, combined with their role in interference competition, could also point towards their role in bacterial speciation. The first evidence for this hypothesis has recently begun to emerge from analysis of closely related, strains^59–61^.

Recent work on virome transplants indicate the tremendous role of phages in human health. It remains to be discovered to what extent this can be assigned to pBGCs, likely due to fitness effects, an easy path of HGT of those BGCs. Finally, prophage activation may occur during antibiotic therapy or even due to diet choices^63^, promoting spread of pBGCs across closely-related strains. Phages represent a treasure chest of new BGCs with new activities. Here, we demonstrated the natural role of these phage-encoded BGCs in interbacterial warfare and prove the plasticity of their transfer to new bacterial host, which can take us forward in application of specialized metabolites. In summary, encoding of bacteriocins could be an evolutionary strategy adapted by some phages to weaponize their host, thereby conferring a competitive advantage of both the host and hence the phage infecting it.

## Materials and methods

### Strains and growth conditions

All bacterial strains used in this study as well as strains used as genomic DNA donors are described in Supplementary table 1. Strains were routinely maintained in lysogeny broth (LB) medium (LB-Lennox, Carl Roth; 10 g l^-1^ tryptone, 5 g l^-1^ yeast extract, and 5 g l^-1^ NaCl). Strain DTUB231 (P9_B1^SPβ^) was obtained by infecting P9_B1 with a *Bacillus* phage SPβ isolated from strain MB8_B7. Isogenic lysogeny was confirmed by colony PCR using primers pairs TB122/oAD2 (Supp. table 1) and oAD28/oAD3 (Supp. table 1) binding to flanking prophage regions and host chromosome close to *attL* or *attR*, respectively. Colonies that were positive for oTB122/oAD2 and oAD28/oAD3, but negative for oAD2/oAD3 (intact *att*) were selected as stable SPβ lysogens. Strain DTUB208 was obtained by transforming MB8_B7 with gDNA of SPmini. DTUB43 and DTUB222 were obtained by transforming P9_B1 with phy_sGFP and phy_mKATE2 plasmids, respectively. Strains DTUB235 and DTUB236 emerged from infecting of DTUB43 and DTUB222 with SPβ phage, respectively. Stable lysogens were confirmed by colony PCR like described above. Strains DTUB233 and DTUB234 were obtained by transforming DTUB43 and DTUB222 with gDNA isolated from GM3248 and selecting for kanamycin-resistant colonies. Strains DTUB244 and DTUB245 were obtained by transforming DTUB43 and DTUB222 with gDNA isolated from Δ*sunA* and selecting for kanamycin-resistant colonies, while DTUB246 and DTUB247 were obtained by transforming DTUB43 and DTUB222 with gDNA isolated from ANC3, also selecting for kanamycin resistance. Deletions were confirmed using oAD61/oAD62 (Supp. table 1) binding to *sunA*-flanking regions. Working concentrations of antibiotics were 5 µg ml^-1^ for erythromycin, kanamycin and chloramphenicol, and 100 µg ml^-1^ for spectinomycin. Primers 27F/1492R (Supp. table 1) targeting 16S rRNA were used as a positive PCR control.

### Phage isolation and purification

Strain MB8_B7 was cultivated at 37 °C with shaking at 200 r.p.m. In mid-exponential phase Mitomycin C was added (1.5 µg ml^-1^) to trigger prophage induction, following cultivation for another 6h. Cells were pelleted down (8000 g, 10 min), supernatant was filtered sterilized and diluted in order to obtain well-separated single plaques. Plaque assays were performed using Δ6 strain as a host. Plaque giving PCR product for SPβ-specific primer pairs oTB88/oTB89 (Supp. table 1) and oAD51/oAD52 (Supp. table 1), was carefully removed from the soft agar using a sterile scalpel, resuspended in 200μl of SM buffer and used to infect exponentially growing phage-free host Δ6 to allow propagation. Phage was subsequently propagated in soft agar and liquid host suspension until the titer reached at least 10^9^ pfu/ml. Such culture supernatants were collected, adjusted to pH of 7.0, filter -sterilized and mixed at a 1:4 rate with PEG-8000 solution (PEG-8000 20 %, 116 g l^-1^ NaCl). After overnight incubation at 4°C, the solutions were centrifuged for 60 min at 12000 rpm to obtain phage precipitates. The pellets were resuspended in 1 % of the initial volume in SM buffer (5.8 g l^-1^ NaCl, 0.96 g l^-1^ MgSO_4_, 6 g l^-1^ Tris-HCl, pH 7.5) to obtain concentrated solution of phage particles.

### Sublancin activity assay

Sublancin activity assays were performed as previously described^34^. Briefly, overnight cultures of selected strains were diluted 100×, and 100 µl of the diluted cultures were transferred onto the LB-agar (1.5 %) plate using a plastic spreader to serve as a lawn (target strain). For the focal strains, undiluted overnight cultures were spotted (5 µl) on top of the lawn. Plates were incubated at 37 °C for 24 h. Clearing zone around the focal colonies indicated their antimicrobial activity towards the target strain.

### Competition assays

Prior to competition assays the potential fitness costs of introducing fluorescent reporters, were examined and it was noted that mKate-labelled strains may have a slight disadvantage in competition (Suppl. Fig. 3). Therefore, all competition assays were performed with controls, where fluorescent reporters were swapped. As the increase in GFP and mKate fluorescence in monoculture matched the growth pattern of corresponding strains (Suppl., Fig.3), they were used as relative strain ratio indicators in co-cultures. In co-cultures compensation correction was applied, to subtract the effect of GFP background (0.044 %) in mKate channel.

Overnight cultures of selected strains were obtained, optical density was measured and cultures were pelleted down (8000 g, 5 min) and re-suspended in 0.9 % NaCl, to reach optical density of 5. Next, 1:1 co-cultures were created by mixing equal volumes of selected strains suspensions. Such co-cultures were then inoculated at 1 % into 200 µl of LB distributed in 96-well microtiter plates. Cultivation was performed in Synergy XHT multi-mode reader (Biotek Instruments, Winooski, VT, US), at 37 °C with linear continuous shaking (3 mm), monitoring the optical density (600 nm) as well as GFP (Ex: 482/20; Em:528/20; Gain: 35) and mKate (Ex: 590/20; Em: 635/32; Gain: 35) fluorescence every 10 min. For long-term competition assay, co-cultures were transferred to fresh LB medium (2.5 % inoculum), every 24 h.

Growth rates were calculated by linear regression on log transformed OD-values in exponential phase.

### Sample Preparation and LC-MS Analysis of Liquid Cultures

To a 15 mL falcon tube magnesium sulfate (280 mg) and sodium chloride (140 mg) was added. Acidified (0.1 % trifluoroacetic acid) acetonitrile (7 ml) was added, and shaken vigorously by hand for a short time. Culture media (7 ml) was then added to the solution. The mixture was extracted by shaking for 30 min at room temperature. The sample was centrifuged (4200 CFU, 5 min) to allow two liquid phases to form, an organic, acetonitrile-rich, upper phase, and an aqueous, salt-rich, lower phase. The aqueous phase was removed and partially evaporated to approximately 5.5 ml. The partially evaporated aqueous phase was then separated by solid phase extract (SPE), as follows: the aqueous extract was loaded onto a reversed-phase SPE cartridge (Phenomenex Strata-X: 3 ml, 30 mg) that had previously been conditioned with methanol (3 ml) and equilibrated with 0.1 % formic acid in H_2_O (2 ml). After loading the sample onto the column, the column was washed with acidified water (0.1 % formic acid, 2 ml), and the analytes were eluted with 20, 40, and 80 % (v/v) acetonitrile containing 0.1 % formic acid (v/v) (1 ml fractions). Each of the SPE eluents were transferred to individual HPLC vials for analysis by LC-MS using 15 µl injection volumes. LC-MS analysis was undertaken on a UHPLC-QTOF system, using gradient elution with reversed phase chromatography. Refer to Supplementary methods for complete details on chemicals and LC-MS conditions. Sublancin chemical features were detected in the 40 % methanol fractions as [M+3H]^3+^ and [M+4H]^4+^ pseudo-molecular ions, as well as a characteristic [M-C_6_O_5_H_10_+4H]^4+^ fragmentation ion. All cultures were extracted and analyzed in triplicate.

### Transmission electron microscopy

Before use, 400 mesh nickel grids with a 3-4 nm thick carbon film, CF400-Ni-UL EMS (Electron Microscopy Sciences, Hatfield, PA, USA) were hydrophilized by 30 s of electric glow discharging. Next, 5 μl of purified phage solutions were applied onto the grids and allowed to adsorb for 1 min. The grids were rinsed 3 times on droplets of milliQ water and subjected to staining with 2 % uranyl acetate. Specifically, with a help of EM grid-grade tweezers, the grids were placed sequentially on droplets of 2 % uranyl acetate solution for 10 s, 2 s and 20 s. Excess uranyl acetate was wicked away using filter paper and the grids were allowed to dry overnight and stored in a desiccator until analysis. Transmission electron microscopy was performed utilizing a FEI Tecnai T12 Biotwin TEM (Thermo Fisher Scientific, Hillsboro, OR, USA) operating at 120 kV located at the Center for Electron Nanoscopy at the Technical University of Denmark, and images were acquired using a Bottom mounted CCD, Gatan Orius SC1000WC (Gatan Inc., Pleasanton, CA, USA).

### pBGC mining

All complete bacterial refseq genomes were downloaded using ncbi-genome-download (https://github.com/kblin/ncbi-genome-download). Next, all genomes were profiled using a custom script in which prophages were found by using ProphET^37^, in which blastp was substituted for DIAMOND for performance reasons. Next, total genomic biosynthetic gene clusters (gBGCs) were found using antiSMASH4^38^ with the –minimal option enabled, again for performance reasons. Third, prophage encoded BGCs (pBGCs) were found by rerunning antiSMASH4 on the predicted prophages. The complete code is available on request.

Certain species are overrepresented in the genome database and many of the genomes are highly similar which may give both additional false positives and negatives. Dereplication of 15,000+ genomes was considered unfeasible.

### Profiling of pBGCs

pBGCs were annotated using blastp against the BACTIBASE database^64^ with default settings. Presence of pBGC core genes outside the prophage regions were investigated by using blastn against the host and non-host genomes with identity of 70 % and a coverage of 50 %.

### Phylogenetic comparisons

Phages positive for pBGCs were further analyzed by average nucleotide identity using pyani (https://github.com/widdowquinn/pyani), and the resulting nucleotide identity matrix was clustered by hierarchical clustering using hclust() from R on the corresponding distance matrix.

A network of pBGCs was made by considering the ANI as an adjacency matrix using the igraph package in R and building a weighted and directed graph.

The Phaster software^65,66^ was used to detect prophage elements in natural *B. subtilis* isolates. Next, the sequence of SPβ-like prophage detected in MB8_B7 was compared with SPβ sequence of *B. subtilis* 168 using blast. Lack of ICEBs1 (or its fragments) in MB8_B7 was confirmed by blast.

### Whole genome sequencing by MiniON Nanopore

Bacterial genomic DNA was isolated using EURex Bacterial and Yeast Genomic DNA Kit, while phage DNA was extracted using the phenol:chloroform method as described previously^67^. The qualities of the extracted gDNA were evaluated by NanoDrop DS-11+ Spectrophotometer (Saveen Werner, Sweden) and electrophoresis, then quantified with the High Sensitivity DNA kit and Qubit 2.0 Fluorometer (Thermo Fisher Scientific). For multiplex MiniON sequencing, the Rapid Barcoding gDNA Sequencing kit was used to allow pooling of samples on a single MinION Flow Cell (FLO-MINSP6). Genomic DNA library was prepared using the SQK-RBK004 rapid barcoding kit and subsequently loaded into the MinION Flow Cell. The local base calling was performed automatically in real-time using the MinKNOW ONT software (v3.1.19). De-multiplexing and adapter trimming was carried out using EPI2ME Agent software (v2020.2.10) using the Fastq Barcoding r2020.03.10 function and the genomes were then assembled using Unicycler v0.4.8^68^.

### Statistical analysis

The association between pBGC positivity versus prophage or gBGC counts was tested for significance using Mann-Whitney U-tests.

To calculate the individual effects of the SPβ prophage, the intact pBGC or only the *sunA* gene on strain performance during competition, the change in proportion from 0h to 24h in a strain with a given gene content was compared to the change in proportion of the non-modified strain with the same fluorescent marker. Statistical differences between two experimental groups were then identified using a two-tailed Student’s *t*-tests assuming equal variance. To assess the effect of phage *att* site (*spsM*) on competition in the presence and absence of SPβ, changes in P9_B1 proportion throughout the competition (start vs 3^rd^ at transfer 24h) in the absence of SPβ, where compared to changes of its proportion during competition with single exposure to SPβ. Relative abundance of strains in co-culture was calculated based on fluorescence values (see Suppl. Fig. 3). Normal distributions within the above datasets were confirmed by Kolmogorov–Smirnov (*P*>0.05). Differences in growth rates between P9_B1, P9_B1_GFP_ and P9_B1_mKate_ were assessed by One-Way ANOVA and mean comparisons by Tukey test. No outliers were removed from the dataset. No statistical methods were used to predetermine sample size and the experiments were not randomized.

## Supporting information

Supplementary Figures + Legends - merged

## Data availability

Raw sequencing data have been deposited to SRA databased under sub8228942 (for ΔspsM^SPβ^) and sub8237787 (for SPβ) accession numbers. The complete code used for mining of phage-encoded BGCs is available on request from Mikael Lenz Strube.

## Authors contributions

A.D. and M.L.S. designed the study. M.L.S. performed bioinformatic analysis, A.D. performed the experiments, A.J.C.A. performed chemical analysis, C.N.L.A. performed Nanopore sequencing and data processing, A.T.K. provided molecular tools for the study. A.D. and M.L.S. wrote the manuscript. All authors contributed to final version of the manuscript.

The authors declare no conflict of interest.

## Acknowledgements

The authors thank L. Gram for feedback during ongoing project and constructive comments on the final version of this manuscript. We also thank A. Kofod-Petersen for valuable discussions. This work was supported by the Danish National Research Foundation (DNRF137) for the Center for Microbial Secondary Metabolites.

