## Supplementary Figures + Legends - merged for "Phages weaponize their bacteria with biosynthetic gene clusters"

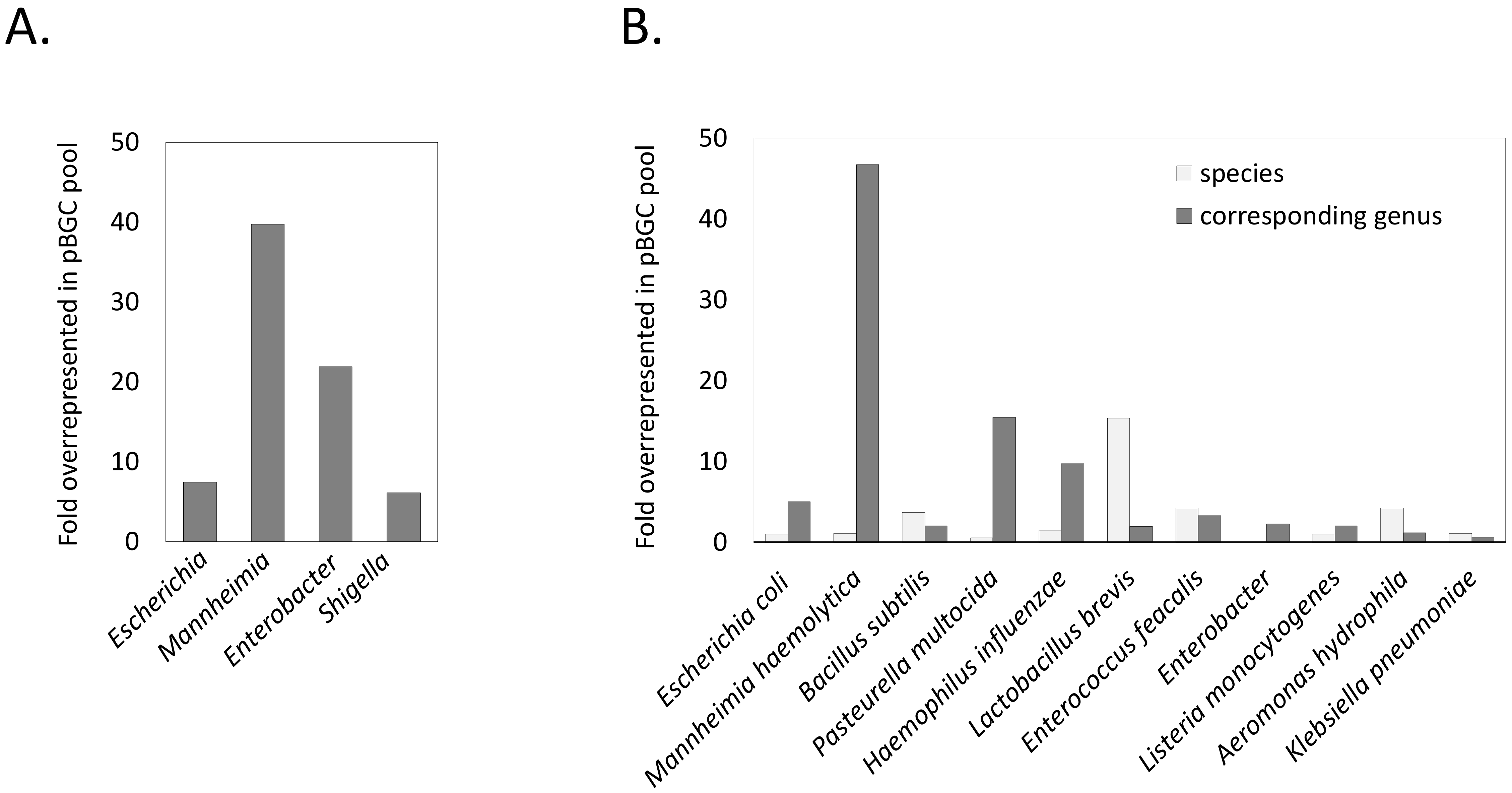


Supp. Fig. 1. Abundance of pBGCs in virions (A) or prophages (B) of different bacterial genera/species, compared to abundance of these genera/species in analyzed databases (PATRIC and NCBI, respectively), represented as fold-overrepresented value - 1 indicates the relative abundance of pBGC per genus/species equals its relative abundance in the analyzed database.

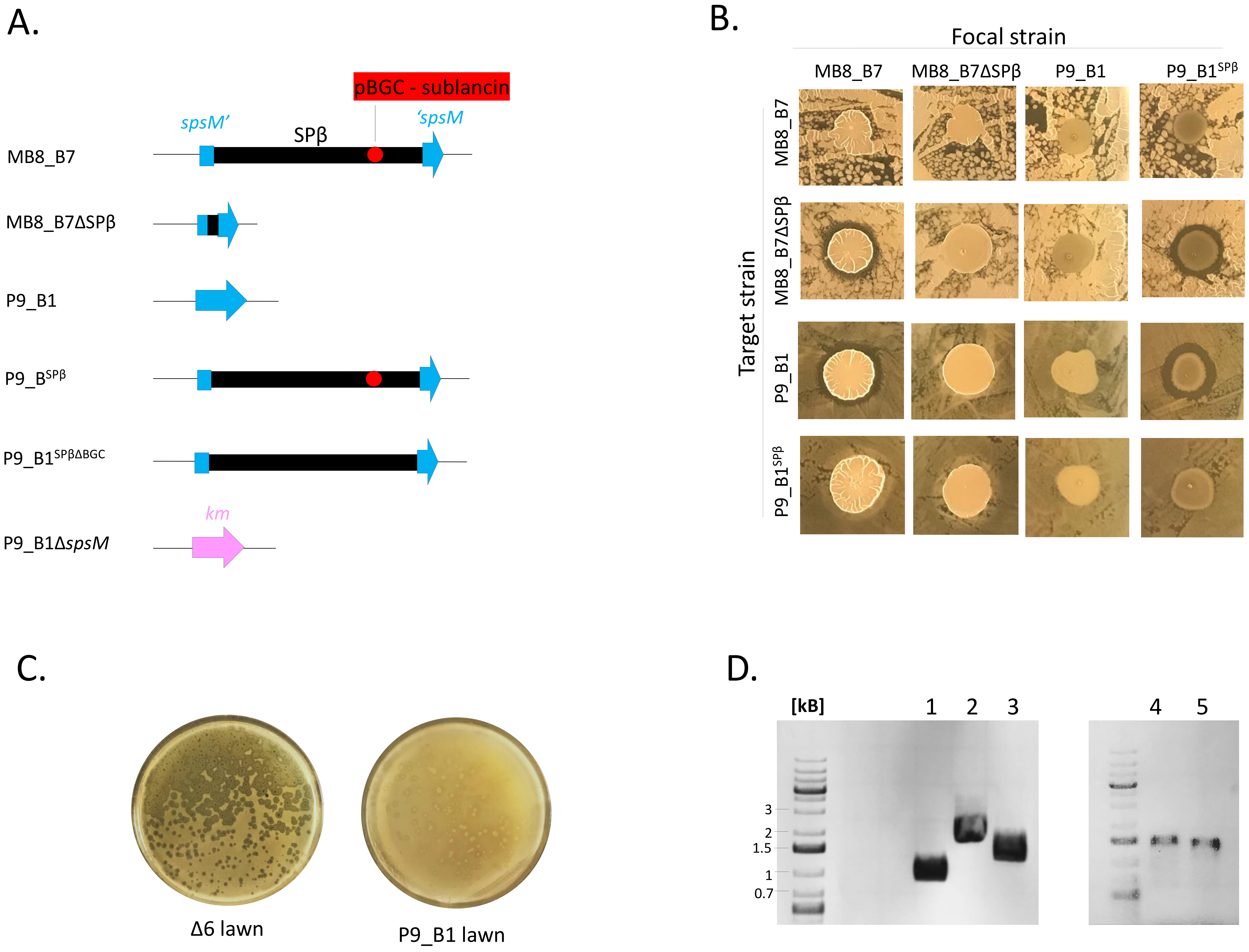


Suppl. Fig. 2. Experimental system to test benefits of a phage-encoded BGC for the host. A. The *Bacillus subtilis* isolate MB8_B7 (CP045822.1) served as a donor of pBGC (sublancin)-carrying phage called SPβ. Gene *spsM* serves as an attachment site for SPβ during lysogenic cycle. The *Bacillus subtilis* isolate P9_B1 (CP045811.1), was selected to test benefits of pBGC – this strain is non-lysogenic for SPβ but carries an intact *spsM* gene potentially allowing the attachment of SPβ and obtaining strains P9_B1$^{SPβ}$ (also referred to as P9_B1$^{SPβ}$) as well as P9_B1$^{SPβΔBGC}$ (also referred to as ΔpBGC). Strain P9_B1Δ*spsM* (also referred to as Δ*spsM*), was used to test benefits from chromosomal attachment site for pBGC-encoding phage. B. Antagonistic assay between MB8_B7, MB8_B7ΔSPβ, P9_B1 and P9_B1 infected with SPβ. Each strain served as a target (when fully grown culture was diluted 1:100 and inoculated as a lawn) and as a focal strain, when fully grown culture (non-diluted) was spotted onto the lawn of the target strain. C. Plaque assay with SPβ obtained from MB8_B7, using the Δ6 strain and P9_B1 strains as a lawn. D. PCR-test confirming SPβ identify after isolation from single plaque, propagation in Δ6 and purification. Primers and expected PCR-product size were as follows: 1 – SPβ region I: oTB88-89/ 1096 bp; 2 – SPβ region II: oAD51-52/ 2096 pb; 3 – PCR positive control on bacterial gDNA: 27F-1492R/ 1473 bp; 4 – SPβ integration, left arm: oTB122-oAD2/ 1149 bp; 5 – SPβ integration, right arm: oAD28-oAD3/ 1141 bp.

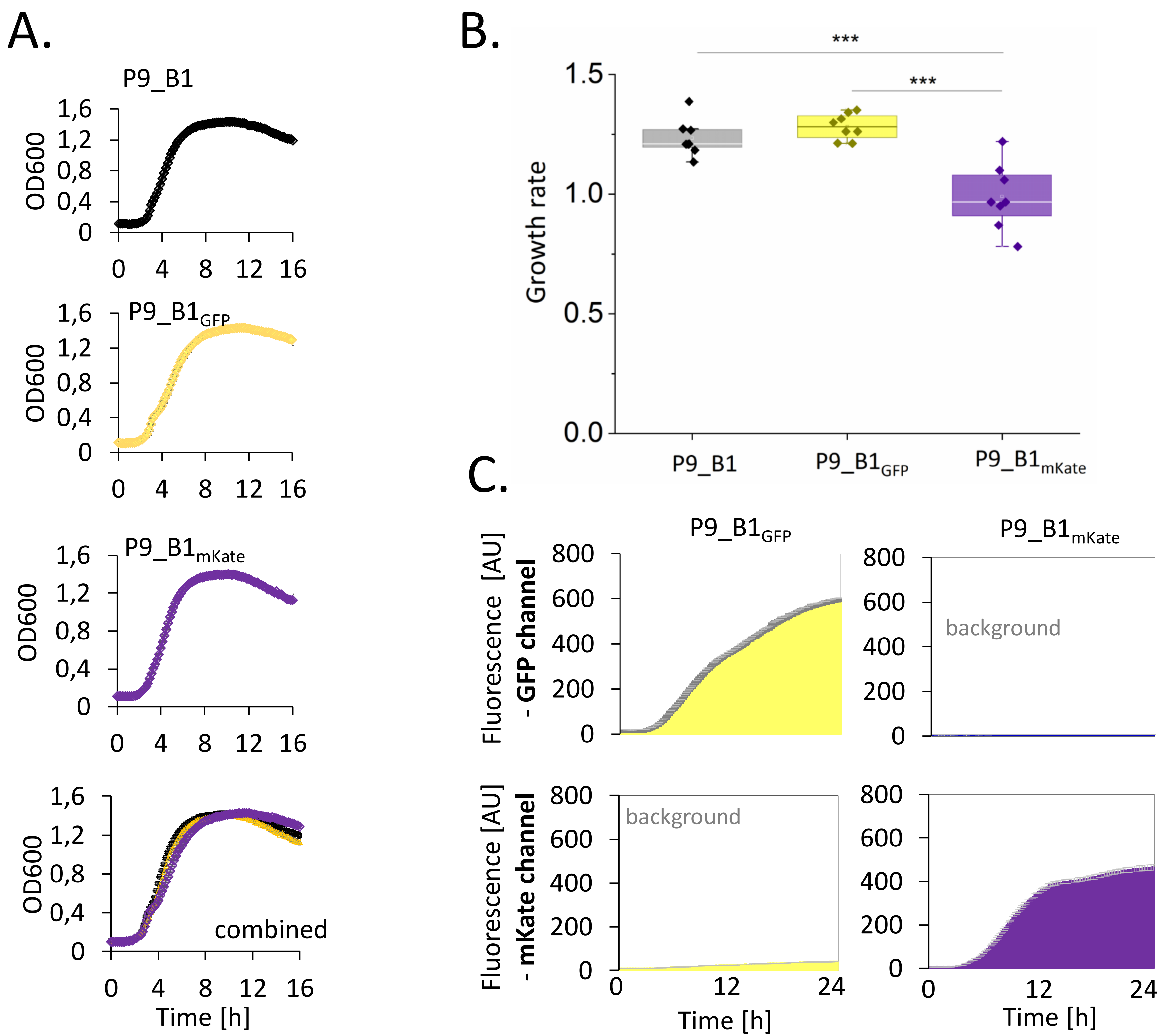


Suppl. Fig. 3. Examination of fluorescently labelled strains used in competition assays. A. Original P9_B1 strain and its fluorescently labelled derivatives were grown in monoculture with monitoring optical density (OD600). B. Growth rates of P9_B1 (non-labelled), P9_B1$_{GFP}$ and P9_B1$_{mKate}$. *** indicates $P<0.001$ ($n=6$). C. Changes of specific and background fluorescence values during growth of P9_B1$_{GFP}$ and P9_B1$_{mKate}$. Data represent average from 6 biological replicates, error bars (light grey) represent standard error.

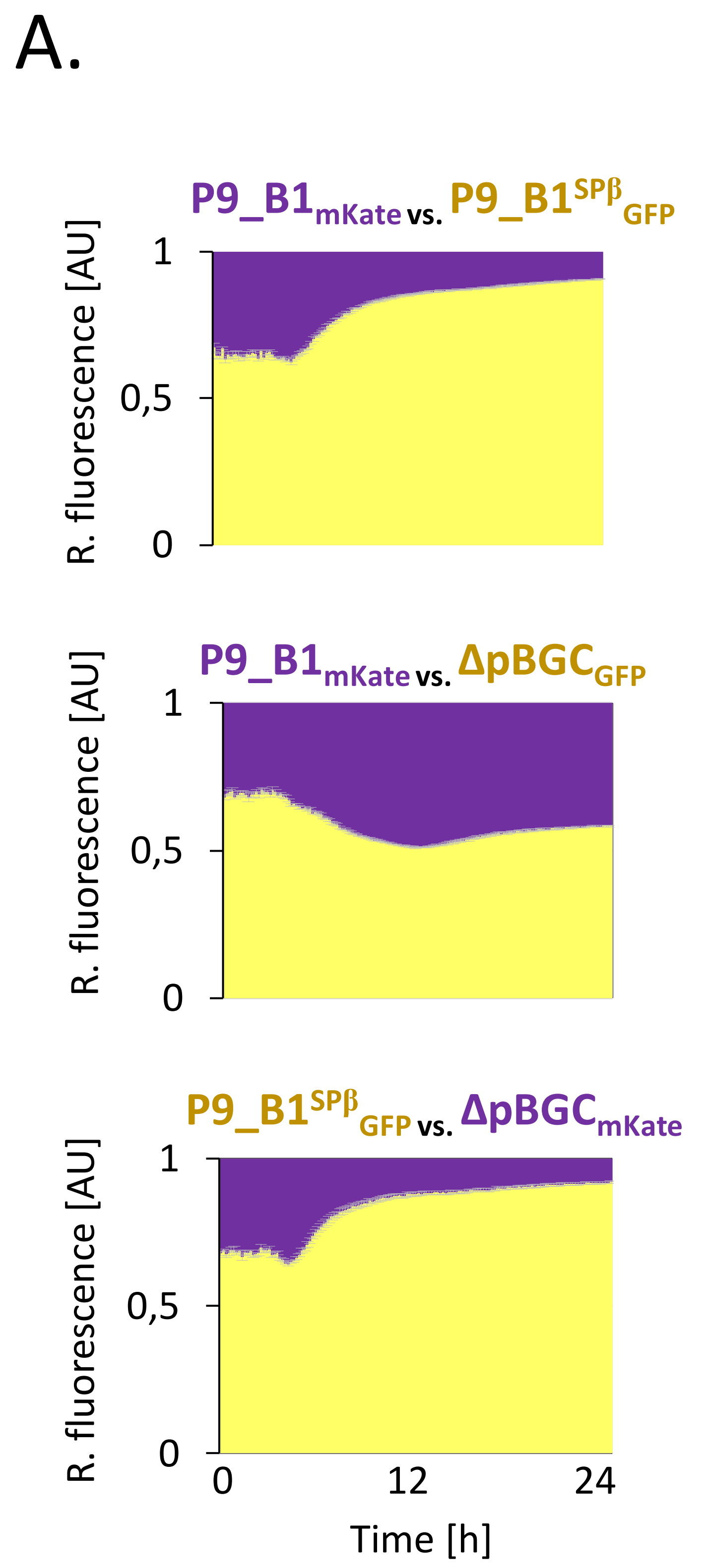


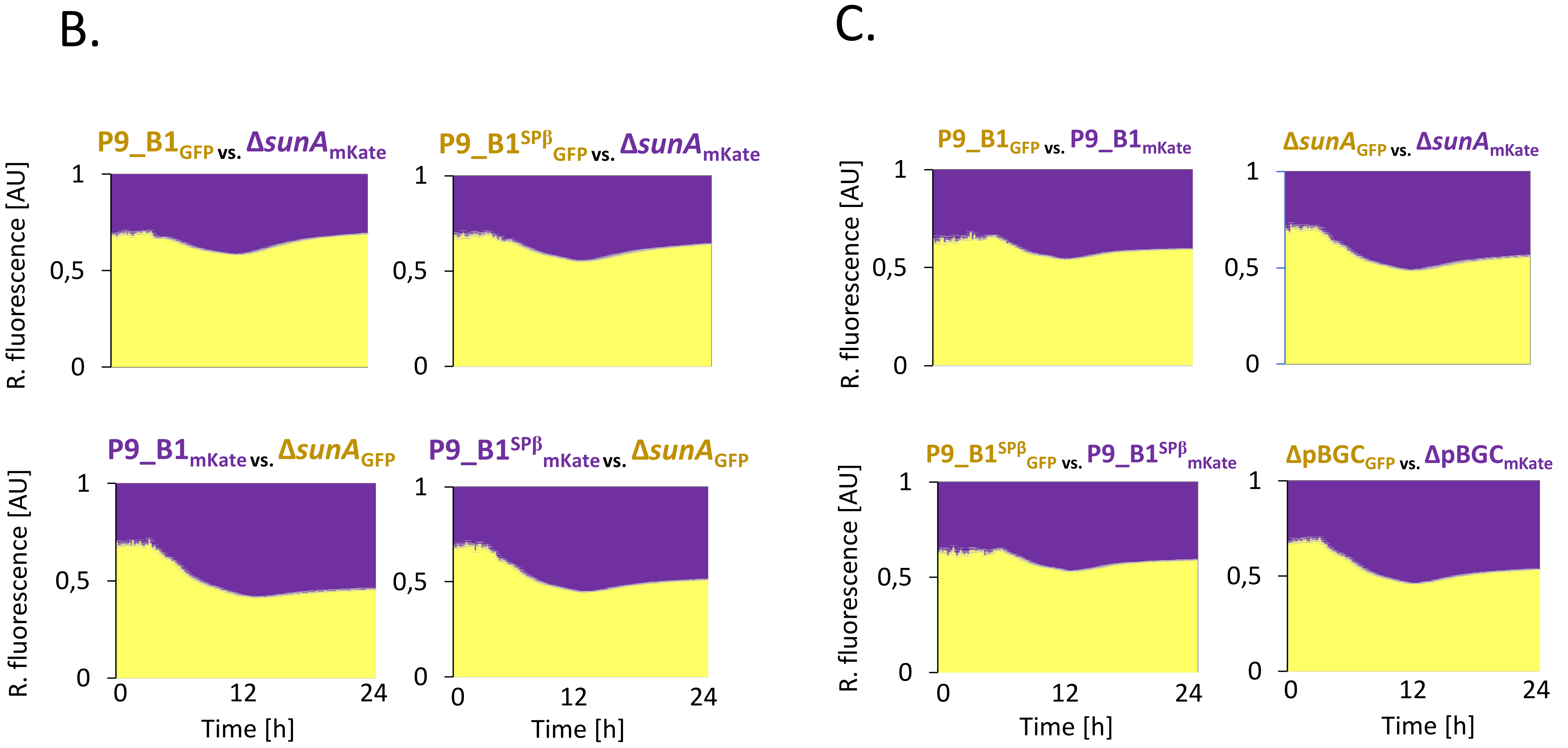


Suppl. Fig. 4. Competition assays between fluorescently labelled strains – control experiments. A. Competition assays involving P9_B1, P9_B1$^{SP\beta}$ and ΔpBGC, swapped fluorescent reporters respect to Fig. 4C. B. Competition assays of Δ*sunA* against the P9_B1 or P9_B1$^{SP\beta}$. C. Competition assays between isogenic strains that differ solely with fluorescent reporters. All assays were initiated from 1:1 ratio of the starter cultures. Relative abundance of two competing strains was monitored by changes in relative GFP and mKate fluorescence, shown in yellow and purple, respectively. Error bars (light grey) indicates standard error (n=6-8).

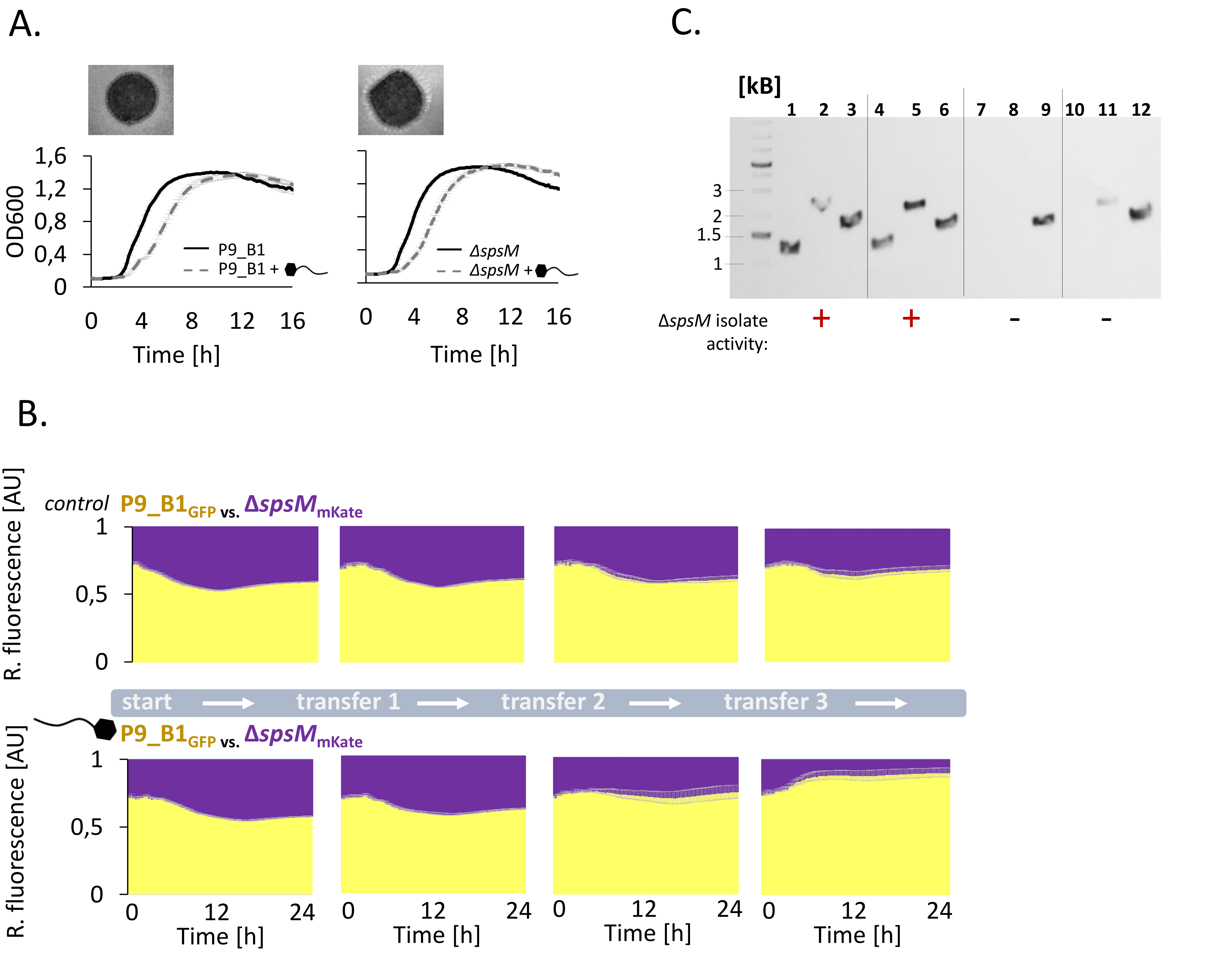


Suppl. Fig. 5. Comparison of P9_B1 and Δ*spsM* – control experiments. A. Growth curve of P9_B1 and Δ*spsM* without and with exposure to SPβ in exponential phase. Above images: spotting of SPβ solution on lawn obtained from exponentially growing P9_B1 and Δ*spsM*. B. Long term competition assay between P9_B1$_{GFP}$ vs Δ*spsM*$_{Kate}$ without and with single exposure to SPβ at the exponential growth phase during first co-cultivation round. All assays were initiated from 1:1. Relative abundance of two competing strains was monitored by changes in relative GFP and mKate fluorescence, shown in yellow and purple, respectively. Error bars (light grey) indicates standard error (n=8). C. PCR screen of Δ*spsM* isolates derived from co-cultures exposed to SPβ. (+) indicates presence of antagonistic activity and (-) indicates absence of antagonistic activity against P9_B1 ancestor. Primers and expected PCR-product size were as follows: 1,4,7,10 - SPβ region I: oTB88-89/ 1096 bp; 2,5,8,11 – SPβ region of pBGC: oAD61-62/ 2001 bp; 3,6,9,12- PCR positive control on bacterial gDNA: 27F-1492R/ 1473 bp.

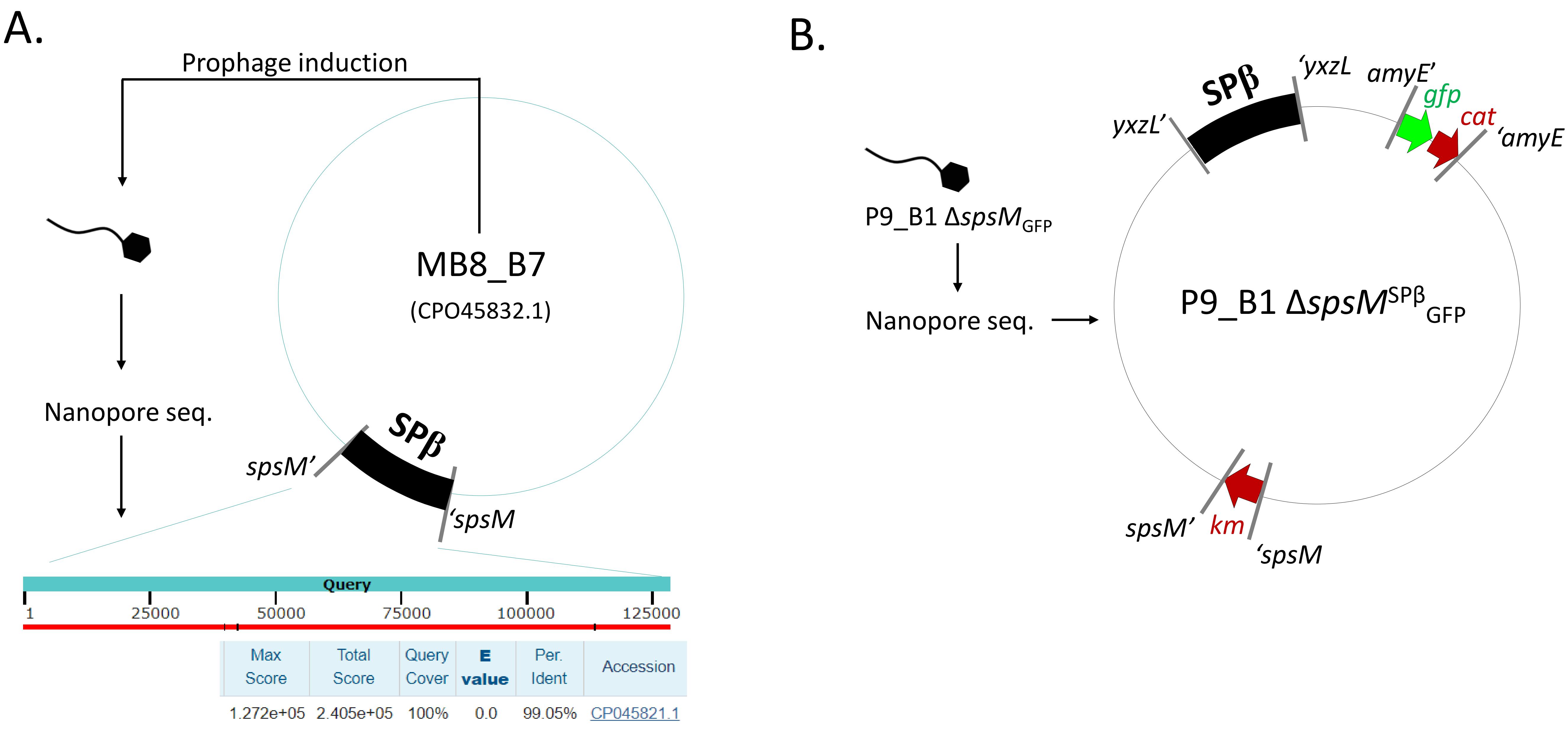


Suppl. Fig. 6. Schematic representation of whole genome sequencing data. Strain MB8_B7 (CP045832.1) was used as SPβ donor. A. Phage DNA was isolated and subjected to Nanopore sequencing (see methods). The resulting sequence obtained by de novo assembly had a 100 % overlap with the SPβ prophage region present in the MB8_B7 chromosome, of which it originated. B. A Δ*spsM* mutants showing antagonistic activity against the P9_B1 ancestor after being exposed to SPβ was subjected to Nanopore sequencing to examine the alternative *att* site for SPβ. Integration of SPβ as well as additional genetic modifications of the strain confirmed by sequencing, are indicated.

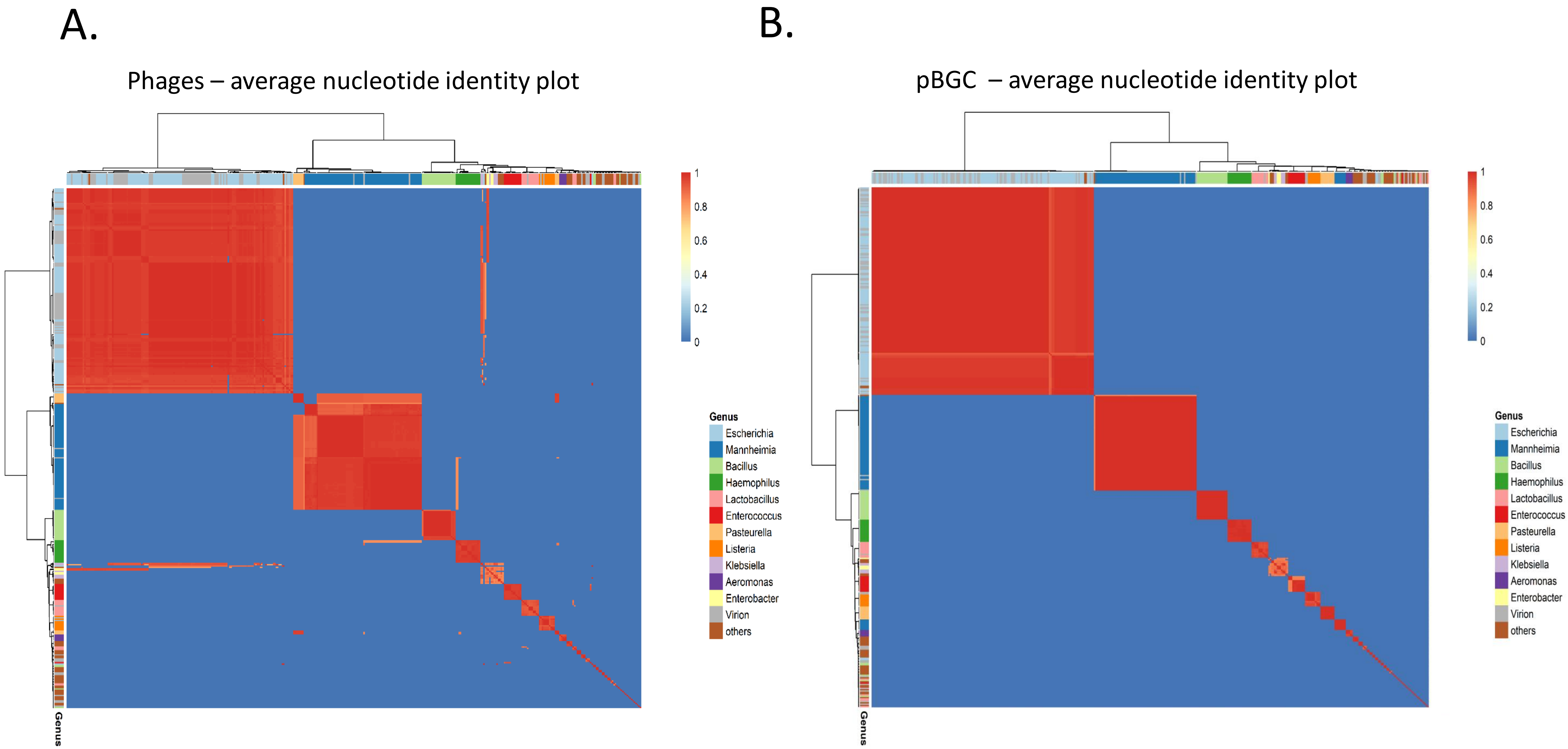


Suppl. Fig. 7. Diversity of phages and pBGCs. A) Average nucleotide identity plots (ANI) of pBGC similarity of full length prophages and virions carrying pBGCs, labelled by major genera. Each edge is weighted by the ANI-value of each pair. B) Network of pBGC core gene similarity of prophage- and virion-derived pBGCs, labelled by major genera. Each edge is weighted by the ANI-value of each pair.
