## Supplementary Table 1 for "Phages weaponize their bacteria with biosynthetic gene clusters"

Suppl. table 1 | Molecular tools used in this study

|  | **Genotype** | **Reference** |
| --- | --- | --- |
| ***Bacterial strains*** | | |
| P9_B1 | Wild type | ^35^ |
| MB8_B7 | Wild type | ^35^ |
| DTUB231 | P9_B1 lysogenic for SPβ phage | This work |
| DTUB208 | MB8_B7 *SPβ::ery* | This work |
| DTUB43 | P9_B1 *amyE::gfp (cm)* | This work |
| DTUB222 | P9_B1 *amyE::mKate (spec)* | This work |
| DTUB235 | P9_B1 *amyE::gfp (cm)* lysogenic for SPβ phage | This work |
| DTUB236 | P9_B1 *amyE::mKate (spec)* lysogenic for SPβ phage | This work |
| DTUB233 | P9_B1 *amyE::gfp (cm), spsM::km* | This work |
| DTUB234 | P9_B1 *amyE::mKate (spec), spsM::km* | This work |
| DTUB244 | P9_B1 *amyE::gfp (cm), sunA::km* | This work |
| DTUB245 | P9_B1 *amyE::mKate (spec), sunA::km* | This work |
| DTUB246 | P9_B1 *amyE::gfp (cm),* yolF, sunA, sunT, bdbA, yolJ, bdbB::km | This work |
| DTUB247 | P9_B1 *amyE::mKate (spec),* yolF, sunA, sunT, bdbA, yolJ, bdbB::km | This work |
| Δ6 | trpC2; ΔSPβ; Δskin; ΔPBSX; Δprophage1; pks::Cm; Δprophage 3 | ^69^ |
| ***Phages*** | | |
| Bacillus phage SPβ | Wild type | This work |
| ***Plasmids*** | | |
| pTB497 | sfGFP with hyspank promoter, Amp (Cm) | ^70^ |
| pTB498 | mKate2 with hyspank promoter, Amp (Spec) | ^71^ |
| ***Bacterial strains used as gDNA donors*** | | |
| SPmini | *SPβ::ery* | ^53^ |
| GM3248 | spsM::km | ^52^ |
| ΔsunA | trpC2, sunA::km | ^34^ |
| ANC3 | trpC2; yolF, sunA, sunT, bdbA, yolJ, bdbB::km | ^34^ |
| ***Primers*** | | |
| oTB122 (F) | TATTGAGCTTGCCAAACTCATAAGAATGAA | |
| oAD2 (R) | CTGCTCTGGAAAGGAAGGCAGAGTAA | |
| oAD28 (F) | GCAGGGCCCCCTACACCGTCAGGGAAA | |
| oAD3 (R) | ATGACCGAACCTCTGGAACCGAGAAC | |
| oAD61 (F) | GCAACATGTGCCTGCTGAAG | |
| oAD62 (R) | GGTATGCCATATGCTCAACC | |
| oTB88 (F) | GATGCCGGTTATCCTTAC | |
| oTB89 (R) | AGGCTAGGCTCTAAGAAG | |
| oAD51 (F) | AGAGCCGGTCAAAGGTAAAC | |
| oAD52 (R) | GCTTTGCTGCAACTGTTG | |
| 27F (F) | AGAGTTTGATCMTGGCTCAG | |
| 1492R (R) | TACGGYTACCTTGTTACGACTT | |
