## Supplementary methods for "Phages weaponize their bacteria with biosynthetic gene clusters"

**Chemicals and LC-MS Analysis**

Chemicals used in sample preparation and LC-MS analysis were as follows. H_2_O used in sample preparation was prepared from a Milli-Q system (Millipore Sigma; Darmstadt, Germany). Acetonitrile (MeCN) and methanol (MeOH) utilized in samples preparation were of HPLC grade and were purchased from VWR (Copenhagen, Denmark). Trifluoroacetic acid (TFA) was 99 % reagent grade, and purchased from MilliporeSigma. H_2_O and MeCN utilized in LC-MS analysis were of LC-MS grade and were purchased from VWR and MilliporeSigma, respectively. Solid phase extraction (SPE) columns were obtained from Phenomenex (Værløse, Denmark). Magnesium sulphate and sodium chloride were purchased from MilliporeSigma. Formic acid was 99 % LC-MS grade and purchased from Fisher Chemicals.

Chromatographic separation was achieved on a Thermo Scientific Dionex UltiMate 3000 RS UHPLC equipped with a Phenomenex F5 column (PFP, 1.7 µm, 100 Å, 150×2.1 mm). Gradient elution was utilized in separation, and the eluents were as follows, Eleunt A: H_2_O, Eleunt B: Acetonitrile. Both eluent A and eluent B contained 20 mM formic acid. The gradient utilized for the chromatographic separation started at 10 % eluent B and increased to 100 % eluent B over 10 min, and was then held at 100 % eluent B for 2.5 min, before returning to the starting conditions. The flow rate and column temperature were kept constant during the entire run, at 400 µl∙min^-1^, and 45 °C, respectively.

The UHPLC was coupled to a Bruker Maxis QTOF. Ionization was undertaken with the use of an electrospray ionization source, and analysis was performed in positive ionization. The *m/z* range selected for detection was 300-2500, and the spectra rate was 2 spectra/sec. The drying gas flow and temperature were set to 10 l∙min^-1^ and 200 °C, respectively. The nebulizer pressure was set to 1.8 bar, and the voltage on the capillary was set to 4500 V. All acquired mass spectra were calibrated with a sodium formate solution
